# Epigenetic responses in *Borrelia*-infected Ixodes *scapularis* ticks: Over-expression of euchromatic histone lysine methyltransferase 2 and no change in DNA methylation

**DOI:** 10.1101/2024.10.25.620284

**Authors:** Grace Hadley MacIntosh, Alexandra C. Nuyens, Jessica L. Vickery, Anne Berthold, Vett K. Lloyd

**Affiliations:** Dept. Biology, Mount Allison University, Sackville, New Brunswick, Canada

**Keywords:** *Ixodes scapularis*, *Borrelia burgdorferi*, Epigenetics, *Euchromatic Histone Lysine Methyltransferase 2*, *G9a*, DNA methylation, *Ixodes Scapularis Repeats*

## Abstract

*Borrelia burgdorferi*, a tick-vectored spirochete bacteria best known for causing Lyme disease, has been found to induce physiological and behavioural changes in its tick vector that can increase tick fitness and its ability to transmit the bacteria. The mechanism by which this bacterium modulates these changes remains unknown. Epigenetics plays a central role in transducing external and internal microbiome environmental influences to the organism, so we investigated DNA methylation and the expression of a key histone modification enzyme in *Borrelia*-infected and uninfected *Ixodes scapularis* ticks. DNA methylation of the pericentromeric tandem repeats family, *Ixodes scapularis* Repeats (ISR) were assessed by methylated-DNA immunoprecipitation (MeDIP) followed by qPCR of the ISR regions. DNA methylation of the ISR sequences was found. The different repeats had different levels of DNA methylation, however, these levels were not significantly affected by the presence or absence of *B. burgdorferi*. The epigenetic regulator *euchromatic histone lysine methyltransferase 2* (*EHMT2*) is recognized as having a key role in modulating the organismal stress response to infections. To assess *EHMT2* transcription in *Borrelia*-infected and uninfected ticks, real-time reverse transcriptase PCR was performed. Uninfected ticks had over 800X lower *EHMT2* expression than infected ticks. This study is among the first to identify a gene that may be involved in producing epigenetic differences in ticks depending on infection status and lays the groundwork for future epigenetic studies of *I. scapularis* in response to *B. burgdorferi* as well as other pathogens that these ticks transmit.

## Introduction

Epigenetic responses mediate the interaction between the environment and the genome to promote homeostasis in the face of external stressors. The environment, including the internal and external microbiota, is the classic trigger of epigenetic changes. The stress response is a prime example of a physiological process that allows an organism to respond to stress, and when the stress is relieved, restore homeostasis [1]. Although, best studied in vertebrates, all organisms, including arthropods, are equipped with a stress response system [2]. Epigenetic processes have long been known to potentiate stress responses by influencing cellular physiology, organismal physiology, differentiation, morphogenesis, variability, both within the lifetime of an organism and between generations, aiding in organismal adaptability [3], [4].

The microbiome is able to modulate multiple physiological functions through both direct metabolite influence on host gene expression and indirectly by epigenetic modulation [5]. Classical examples are baculovirus-mediated control of caterpillar behavior [6], *Ophiocordyceps* manipulation of ant behavior [7], [8] and the complex manipulation of host reproduction by *Wolbachia* spp. bacteria [9], [10]. These studies demonstrate the influence of microorganisms on arthropod behaviors, often by inducing epigenetic modifications of the host.

Epigenetic control relies on complex inter-dependent processes of histone modification, and DNA modification, orchestrated by non-coding RNAs [3], [4], [11], [12]. The *euchromatic histone lysine methyltransferase 2* (*EHMT2*) gene is recognized as a master epigenetic regulatory gene that is involved in multiple biological processes, including restoring homeostasis following physiological stress [1], [13], [14], [15]. Mammals have two subunits: GLP and G9a, whereas invertebrates have been found to have only one [16]. *EHMT*s are characterized by their SET domains, domains that typically have methyltransferase activity that tend to promote transcriptional repression through the mono-(H3K9me) and dimethylation (H3K9me2) of histone H3 at H3K9 and H3K27 [15], [16]. In *Drosophila, EHMT2* promotes survival when the organism is subjected to oxidative stress or viral infection, positioning *EHMT2* as a key epigenetic regulator of stress responses [13], [14]. Interestingly, *EHMT2* has also been shown to have an analogous role in mammalian stress responses by modulating the interferon-mediated immune response in mice embryonic fibroblasts [1], [17]. *EHMT2* expression attenuates the stress response pathways, allowing pathogen tolerance that prevents succumbing to the stressor due to overconsumption of energy and resources [13], [14].

Ticks are hematophagous arthropods and are the vectors of the greatest diversity of pathogens among arthropod vectors, indicating appreciable tolerance for diverse microbiota [18]. Ticks vector multiple pathogens, including *Borrelia burgdorferi*, a prominent pathogen for human and veterinary health [18], [19], [20], [21], [22]. Ticks acquire pathogens upon blood feeding from an infected animal, and then may transmit the pathogen to other hosts upon their next blood meal [23]. Once *B. burgdorferi* is transmitted to non-adapted mammals such as humans or dogs, it can cause Lyme disease. As the spirochetes disseminates throughout the host’s body, it can have far-reaching effects with a broad range of multi-system clinical symptoms [24], [25]. The manifold effects of climate change affect interactions between the environment, animals and humans. Ectothermic arthropod vectors such as ticks are particularly influenced, as are the distribution of their reservoir hosts, all leading to increased infections of humans, companion and agricultural animals [26], [27], [28], [29]; human sero-reactivity to Borrelia has been estimated to be approximately 14.5%, depending on location, detection methodology and risk factors [30].

*Borrelia burgdorferi* and related members of the *Borrelia* genus have adapted to use ticks as vectors [18], [31], [32]. *Borrelia* spirochetes are able to adapt to the dramatically different environments of the arthropod vector and the vertebrate host by sophisticated regulation of its own gene expression [18] as well as modulating the gene expression of its vertebrate host to promote its survival [18], [19], [33], [34]. In the host, changes in gene expression are often seen in genes related to the extra-cellular matrix, the stress response, actin interactions, and the immune response, increasing the probability of the pathogen establishing in the host [34]. Manipulation of its tick vectors is much less studied. As the vector of the pathogen, natural selection would act against a deleterious effect of *Borrelia* on the tick. That consideration does not, however, preclude other changes to gene expression in the vector, in particular those that promote pathogen maintenance in the tick, tick fitness and transfer to the host.

Physiological, behavioral and underlying genetic changes have been detected in *Ixodes ricinus* species complex ticks [18], [20] of which *Ixodes scapularis* and *I. persulcatus* are members. *I. scapularis* is the most common member of this group in North America with a genome 89% homologous to *I. ricinus* [35]. Studies on *I. ricinus* found that *B. burgdorferi* presence in the tick was correlated with increased tick fitness [36]. Ticks infected with *Borrelia spp*. had a greater reserve of fat stores than those uninfected, which reduced mortality from desiccation. Furthermore, infected ticks also showed reduced locomotion, thus conserving energy [37]. Similar complex life-stage dependent changes in questing behavior have been documented in *I. persulcatus* [38] and *I. scapularis* [39]. Although these physiological changes did not correlate with increased pathogen prevalence in one natural setting [40], the lag between tick population establishment and pathogen prevalence [41] would be a plausible explanation for this discrepancy. These physiological changes are underpinned by changes in the expression of genes responsible for nutrient accumulation, lipid production, carbon use, and metabolism, the expression of which are altered in *Borrelia*-infected ticks compared with uninfected ticks [42]. One clear example of manipulation of the tick genome is evidenced by the tick-encoded protein TROSPA. This receptor binds *B. burgdorferi* outer surface protein A (OspA) to allow the bacteria to colonize the tick. The receptor is more highly expressed in *Borrelia*-infected than uninfected ticks, indicating control of tick gene expression to the benefit of the bacteria [43]. *Borrelia burgdorferi* manipulates the expression of other tick genes to enhance vector fitness. In *I. scapularis, B. burgdorferi* infection correlates with increased expression of a tick salivary protein, tick histamine release factor, which leads to enhanced tick feeding, and potentially also enhanced *B. burgdorferi* transmission [44]. *Borrelia burgdorferi* also appears able to manipulate the tick vector to aid the bacteria in establishing infection in the host. Tick salivary proteins Salp15 binds to the *Borrelia* protein OspC when the bacteria is introduced into the host and protects the pathogen from the host immune system [18], [45], [46]. These gene expression changes reflect changes in tick gene transcription coupled to the presence of *Borrelia*, suggesting multiple manipulations of the vector genome to promote pathogen transmission.

These physiological changes and the underlying changes in gene expression promote increase tick fitness, and so foster improved transmission of the *Borrelia* bacteria. However, the underlying epigenetic mechanisms of this process remain largely lacking for *Borrelia*. In contrast, epigenetic control of the tick vector by *Anaplasma* and related tick-borne intracellular bacteria has been investigated [47], [48], [49]. Selection for vector manipulation in obligate tick-transmitted bacteria would be expected to be strong. Thus, demonstrating an epigenetic change in response to *B. burgdorferi* infection is an important first step in a more complete understanding of how tick pathogens modulate physiological changes in their vector to increase the likelihood of pathogen transmission.

DNA methylation has been identified in diverse arachnids, including ticks, although it has been found to be reduced in many parisitiforms [51]. Unlike classical vertebrate systems, DNA methylation tends to be found in gene bodies rather than regulatory regions and may serve to promote rather than repress transcription [51], [52]. As is true of vertebrates, the extent, targets and, to some extent, mechanism of DNA methylation, varies between species and the variability of methylation in invertebrates suggests that it also serves non-transcriptional functions [53], [54]. *I. ricinus* has transcripts encoding DNA methyltransferase (DNMT) enzymes and methylation occurs at the 5-carbon position of cytosine bases [55]. *Ixodes scapularis* has the corresponding DNA sequences and pericentromeric tandem repeats, called *Ixodes scapularis* repeats (ISRs), which were found to be methylated by both methylation-sensitive restriction enzyme digests [56] and binding of antibodies against 5-methylcytosine to tick DNA [55]. These repeats appear to be non-coding and their function is currently unknown [56]. The *I. scapularis* genome has robust representation of families of tandem repeats, constituting 40% of the genome, with ISR-2 repeat family being the most common, constituting approximately 7% of the genome [23]. The limited DNA methylation present in *I. scapularis* may have discouraged research interest into epigenetic regulation in this species. Nevertheless, Nwanade et al. [57] found modest changes in DNA methylation in response to temperature in the Asian longhorned tick, *Haemaphysalis longicornis*, so DNA methylation may be a relevant epigenetic mechanism in ticks.

Histone modifications and non-coding RNA (ncRNA) regulation are the most complex players in epigenetic regulation and orchestrate complex epigenetic responses to the environment in many species, including ticks [20], [49], [58], [59]. The requirement of the histone methyltransferase DOT1L for development in the soft tick *Ornithodoros moubata* exemplifies the essential nature of epigenetic gene regulation [60]. *I. scapularis* has 34 histone modifying enzymes and shows a complex epigenetic response to *A. phagocytophilum* infection [49]. Additionally, Gulia-Nuss et al. [23] identified 4439 ncRNA genes in the *I. scapularis* genome. Although the function of each ncRNA encoded by these genes remains to be elucidated, Artigas-Jerónimo et al. [61] found that *A. phagocytophilum* modulates the miRNA profile of ticks during infection.

The goal of this work was to investigate the epigenetic impact of *B. burgdorferi* on DNA methylation and histone modifications in the tick *I. scapularis*. The epigenetic effects of this pathogen on gene expression in the vertebrate host has been documented, but appreciably less work exists on the pathogen’s effect on the tick vector. Investigation of altered histone modifications in *I. scapularis* in response to *B. burgdorferi* infection is hampered by absence of known gene targets of *Borrelia* genome manipulation. Consequently, we examined expression of the key epigenetic regulator, *EHMT2*. This is a gene with both a well-defined role as an epigenetic regulator and a known role in the stress response and pathogen tolerance, a key physiological response expected in a vector organism.

## Results

### Analysis of DNA methylation of *Ixodes scapularis* centromeric repeat regions (ISR)

For the DNA methylation study, DNA was extracted from 26 female *I. scapularis* ticks, 10 infected with *Borrelia burgdorferi*, 16 uninfected. The concentration of the extracted DNA and 260/280, and 260/230 ratios of archived tick bank samples were assessed and were considered in selecting which samples to include in the groups of tested ticks. Ticks infected with *Anaplasma* spp. or *Babesia* spp. (Table 1) were excluded. Following sonication, as described in Materials and Methods, MeDIP was used to isolate methylated DNA and quantified through qPCR targeting the ISR regions. Control qPCR using the meDNA and unDNA primers, which target the methylated or unmethylated spike-in controls, provided with the MagMeDIP qPCR kit were performed to ensure success of the MeDIP in recovering methylated DNA. Supplemental Figure 4 shows the qPCR results for these controls and confirms successful methylated DNA recovery. The relative amount of methylated DNA, normalized to the input DNA, was then calculated as described in section 2.5 of the Materials and Methods.

**Table 1.**
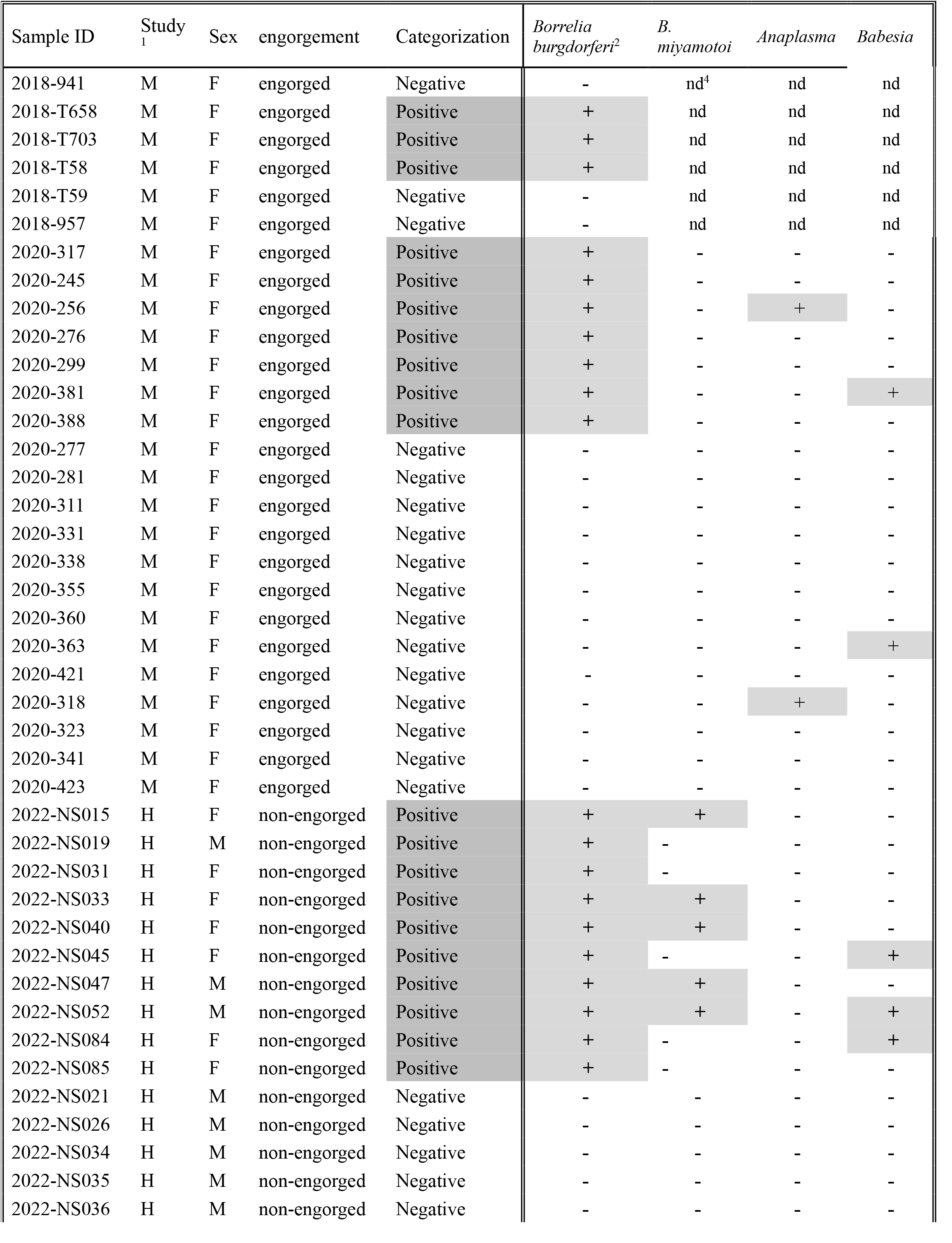
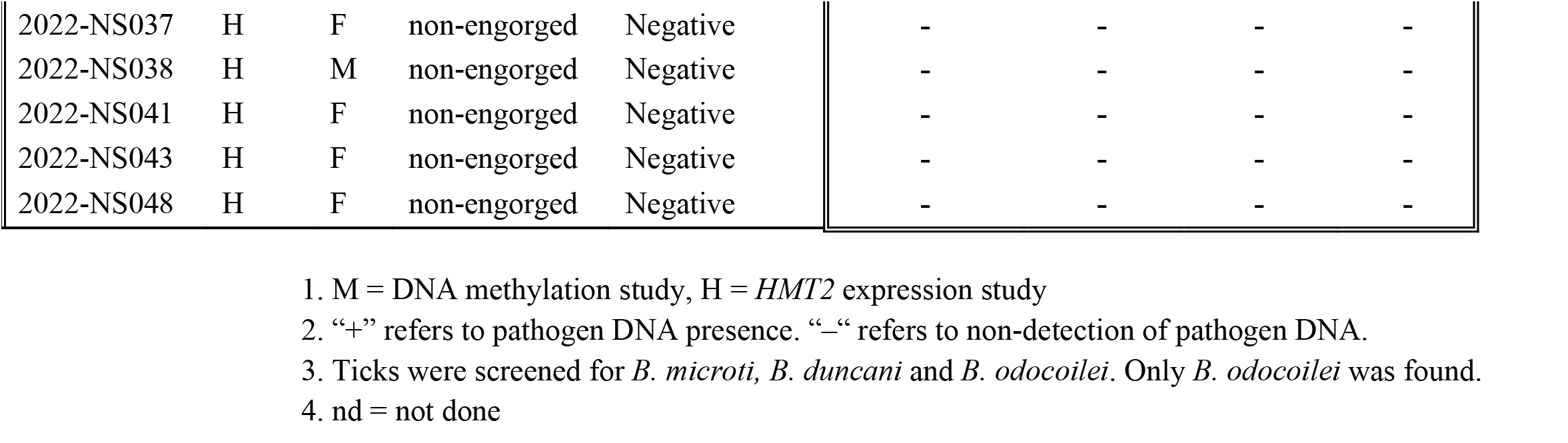
Ticks used to assess epigenetic effects of *Borrelia burgdorferi* on DNA methylation and expression of *EHMT2* in *Ixodes scapularis*.

qPCR was conducted on the methyl-enriched and input DNA samples for each primer region. The PCR amplification curves and melt curves from each *ISR* primer sets is shown in Figure 1. The melt curves show predominantly single peaks, as expected for the amplification of the desired targets, although ISR1 and ISR2C showed multiple peaks, suggesting amplification of multiple complex repeats occurred. The average Ct values and proportion of methylated DNA was calculated from the replicates of each IP and input samples. Figure 2 shows that the relative recovery of methylated DNA from each ISR region, after normalization with the input DNA, for both *Borrelia* positive and negative ticks (Figure 2A) and for the different ISRs without regard to *Borrelia* infection status (Figure 2B). *ISR1* was excluded from this analysis because of erratic amplification precluded generation of a standard curve for the primer set for this region. Not all ticks showed amplification for each primer set so that n= 24 for ISR2A, 20 for ISR2B, 22 for ISER2C, 24 for ISR2D, and 25 for ISR3.

**Figure 1.**
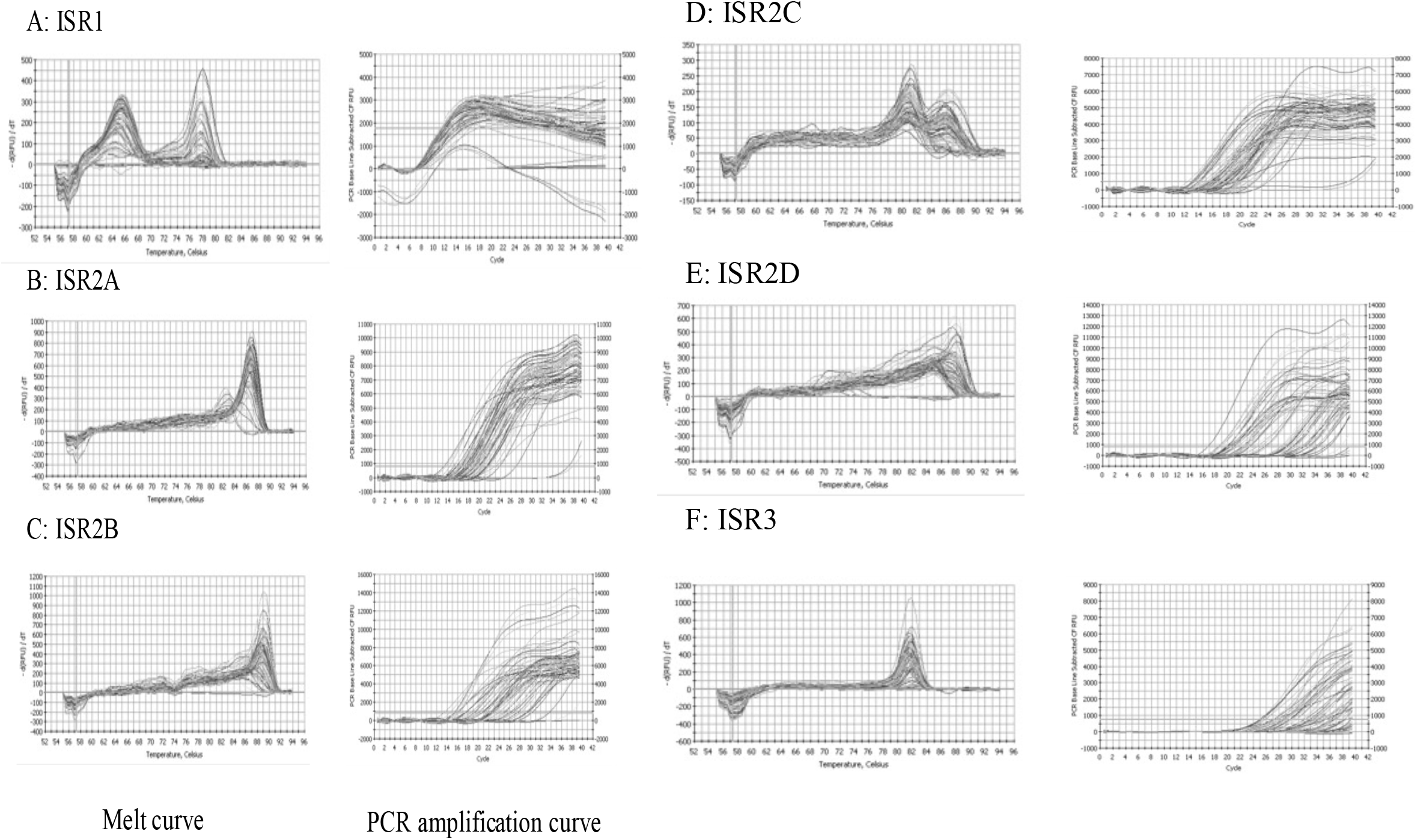
Melt curves and amplification curves for the ISR regions. One peak in the melt curve indicates that one amplicon is being amplified, whereas multiple peaks in the melt curve indicate multiple amplicons. A: Melt curve and amplification curve of ISR1 at 51°C. B: Melt curve and amplification curve of ISR2A at 55°C. C: Melt curve and amplification curve of ISR2B at 55°C. D: Melt curve and amplification curve of ISR2C at 49°C. E: Melt curve and amplification curve of ISR2D at 55°C. F: Melt curve and amplification curve of ISR3 at 55°C.

**Figure 2.**
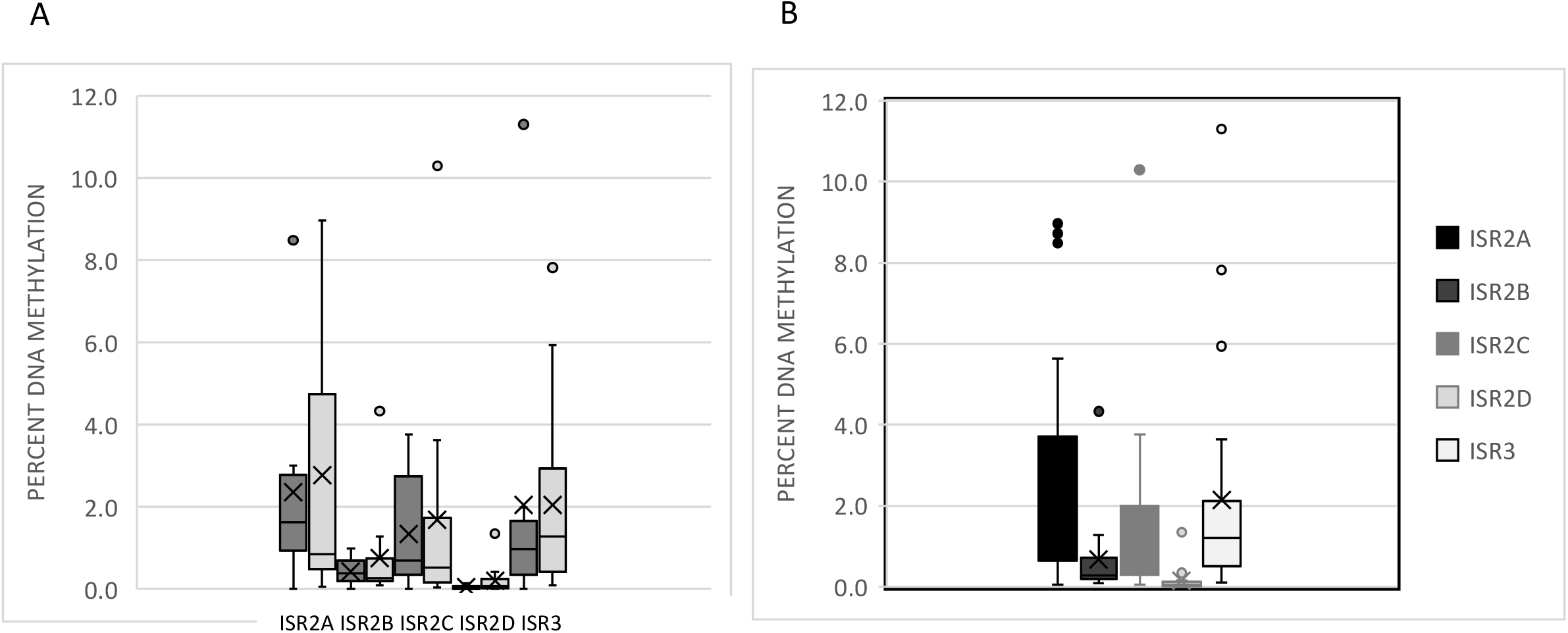
Relative percent DNA methylation of ISR regions ISR2A ISR2B, ISR2C, ISR2D and ISR3. A.ISR regions ISR2A ISR2B, ISR2C, ISR2D and ISR3 (left to right), *Borrelia*-infected (dark grey) and uninfected (light grey). **B**. Relative percent DNA methylation of each ISR region with *Borrelia*-infected and uninfected tick values combined.

Visual inspection of the methylated DNA recovery from *Borrelia*-infected (*Borrelia* (+)) ticks and uninfected (*Borrelia* (-)) (Figure 2A), supported by unpaired t-tests, for each *ISR* region indicates that there was no significant difference between the infection statuses for any of the *ISR* regions (p= 0.91, 0.96, 0.70, 0.93, 0.30, and 0.86 for *ISR1, ISR2A, ISR2B, ISR2C, ISR2D* and *ISR3*, respectively). As there was no significant difference between *Borrelia* (+) and *Borrelia* (-) samples, these values were combined to assess any differences in DNA methylation between the different *ISR* regions. Figure 2B, supported by an ANOVA, shows that the *ISRs* exhibits different levels of DNA methylation with *ISR2A, ISR2C* and *ISR3* showing higher relative levels of DNA methylation than *ISR2B* and *ISR2D*. The *f*-ratio value was 4.56247 and *p*-value 0.002103. Post-hoc Turkey t-tests showed there was a significant difference between *ISR2A* and *ISR2B* (p=0.039), *ISR2A* and *ISR2D* (p=0.004) and *ISR2D* and *ISR3* (p=0.49). These results indicate that the centromeric repeats are methylated and show that the different repeats are methylated to different extents. When normalized to the recovery of methylated DNA, there is no evidence that the presence or absence of *Borrelia* influences the DNA methylation of either highly, or modestly, methylated *ISR* centromeric repeats.

### 3.2 Histone modification – *Euchromatic histone lysine methyltransferase 2 (EHMT2)*

We next sought to investigate whether *Borrelia* presence would affect histone methylation in ticks. In the absence of existing knowledge of which histone modifications might be relevant, rather than targeting specific histone modifications at specific genes, we started by investigating whether *Borrelia* presence altered the transcription of the epigenetic master gene, *EHMT2*. Wild-captured non-engorged ticks were screened for common pathogens and RNA was extracted from 10 of *Borrelia*-infected and 10 uninfected ticks (Table 1, Supplemental Table 2), followed by cDNA synthesis and qPCR for both the *EHMT2* gene and *l13a*, a housekeeping gene as a control gene.

**Table 2.**
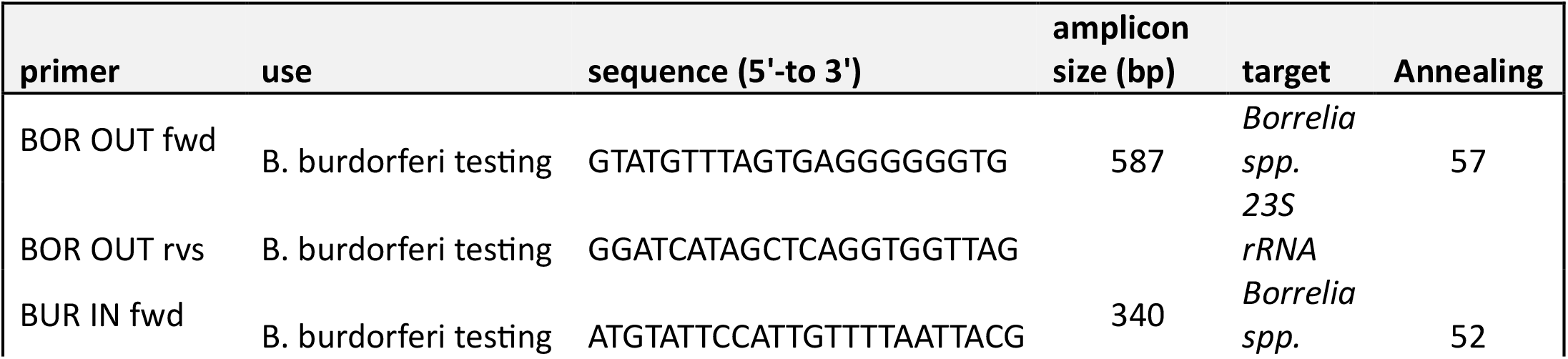
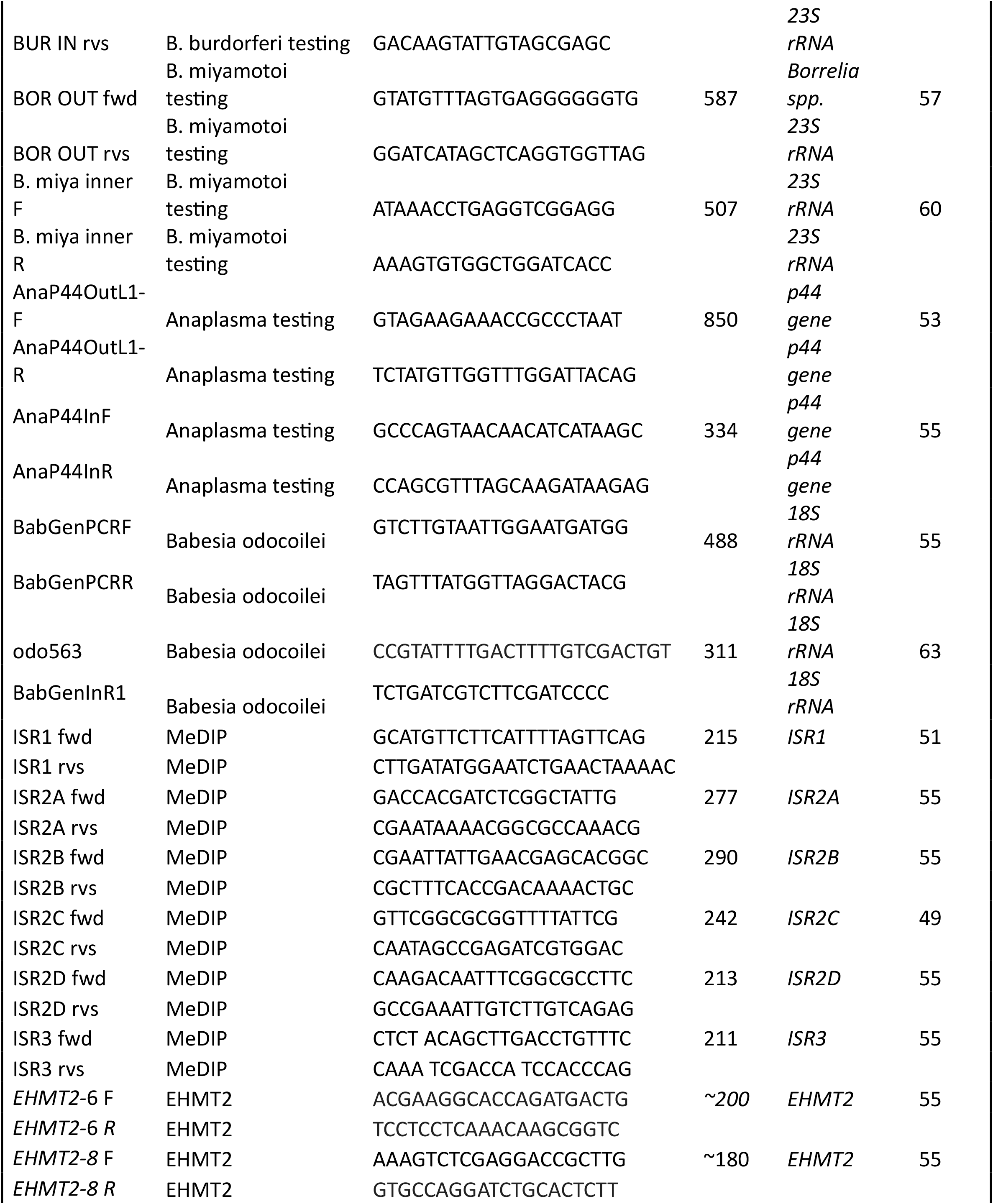
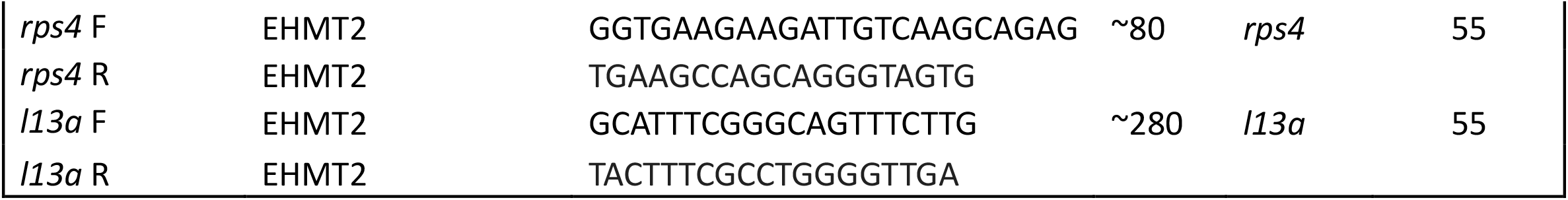
Primers.

Both sets of *EHMT2* primers, *EHMH2*-6 and *EHMH2*-8, and candidate reference genes, *l13a* and *rps4* amplified tick-derived cDNA well, based on both Ct values and confirmatory gel electrophoresis (Supplemental Table 3, Supplemental Figure 5). As described in the Materials and Methods section 2.3.2, the *l13a* primer set was more sensitive than the *rps4* primer set with the *l13a* primers consistently yielding lower Ct values than rps4, so *l13a* was used as the reference gene. The DNase-treated no-RT controls of each tick also underwent qPCR with the *EHMT2*-6, *EHMT2*-8, and *l13a* primer sets. With the exception of one sample, NS038 which was removed from the study, no samples showed amplification, indicating successful cDNA synthesis without contaminating genomic DNA (Supplemental Table 4, Supplemental Figure 6).

**Table 3.**
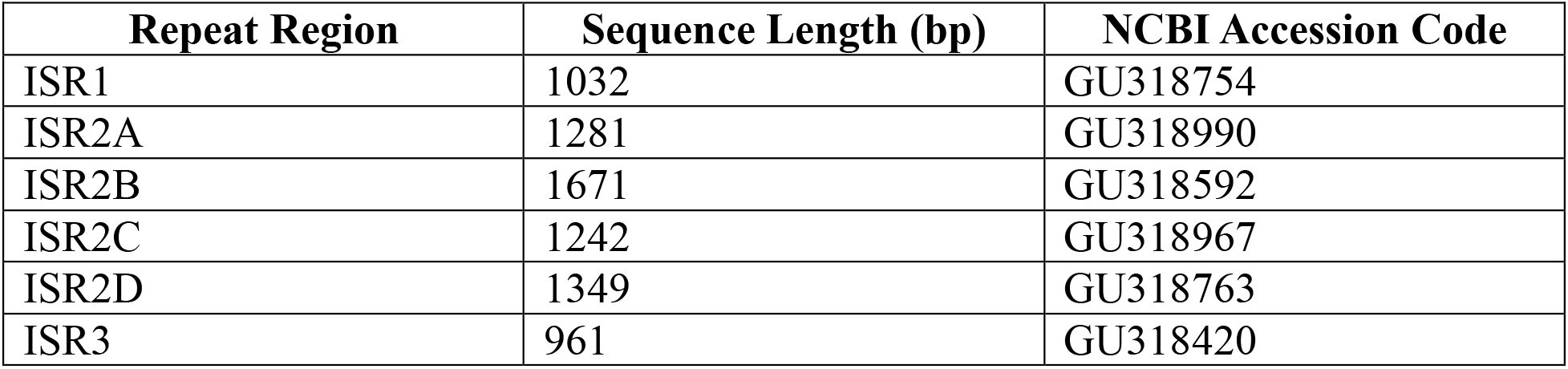
Sequence Length and NCBI Accession Codes for the six *ISR* Repeats.

**Table 4.** Calculation of *EHMT2* expression change in *Borrelia*-infected and uninfected ticks, normalized to the housekeeping gene.

| Designation | TE | TC | HE | HC | ΔCTE | ΔCTC | $\Delta\Delta$Ct | Expression change |
| --- | --- | --- | --- | --- | --- | --- | --- | --- |
| Condition | *Borrelia*-infected | *Borrelia*-uninfected | *Borrelia*-infected | *Borrelia*-uninfected |  |  |  |  |
| Test gene and primer | *EHMT2*-6 | *EHMT2*-6 | *l13a* | *l13a* | TE-HE | TC-HC | ΔCTE-ΔCTC | $2-\Delta\Delta$Ct |
| Average Ct | 34.74 | 33.75 | 33.856 | 23.185 | 0.884 | 10.57 | -9.69 | 826 |

To assess differences between *EHMT2* transcription in infected and uninfected ticks, the Ct values from each tick (excluding NS038) were plotted, along with the Ct values for the control gene, *l13a*. The plot for each *EHMT2* primer set, *EHMT2-*6 and *EHMT2-*8, (Figure 3A and 3B) suggests that the *EHMT2-*6 primers are more sensitive than *EHMT2-*8 primers, as evidenced by lower Ct values and clearer banding when the amplification products are visualized by electrophoresis (Supplemental Figure 7). Thus, the ensuing analysis was conducted with only the *EHMT2-*6 data.

**Figure 3.**
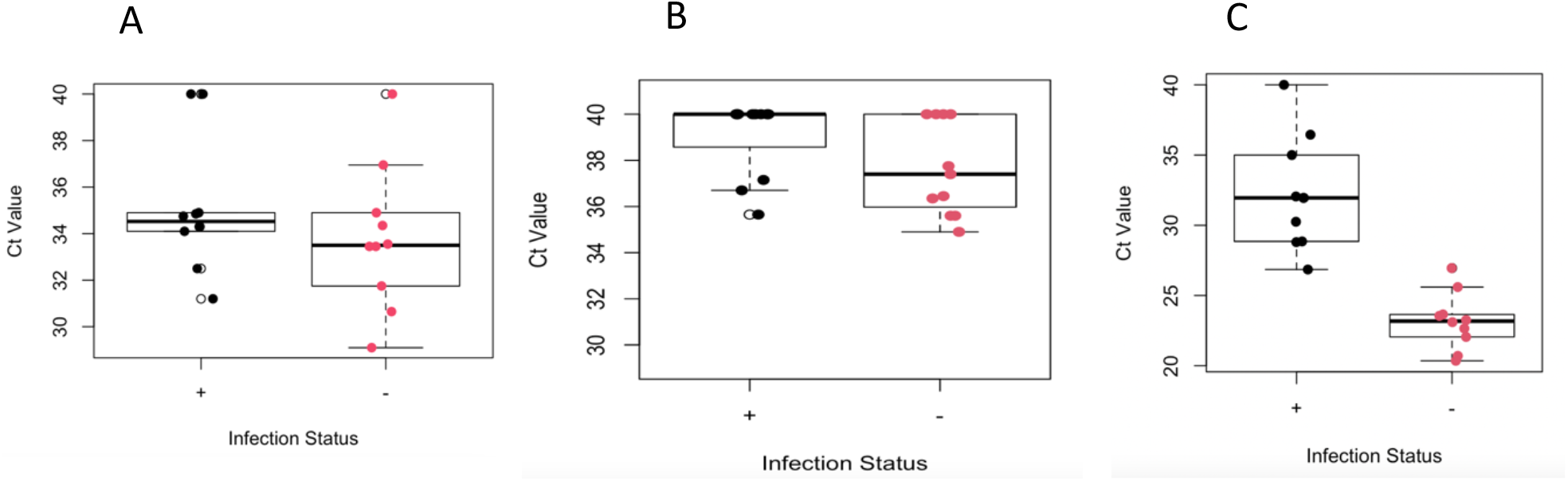
Boxplots of Ct values for *EHMT2* and control gene for ticks with *Borrelia* (black) or without *Borrelia* (red). A) Plot generated with the Ct values produced from qPCR with *EHMT2 6* primers. B) Plot generated with Ct values produced from *EHMT2 8* primers. C) Plot generated with the Ct values with the control gene, *l13a*, primers.

Double delta Ct analysis was used to normalize the *EHMT2* results to the housekeeping gene, *l13a*. This analysis calculates the difference between the expressions of the test gene and the housekeeping gene in each sample, then assesses the difference in that differential between the two experimental conditions – infected and uninfected. The double delta Ct analysis operates under assumptions that there is equal primer efficiency between primer sets; that there is near 100% amplification efficacy of both experimental and control genes; and that the internal control gene, *l13a* in this case, is constantly expressed regardless of treatment [62]. These conditions were met, as described in the Materials and Methods. Figure 3C shows the Ct values for the *l13a* primers. The lower Ct values from the uninfected tick samples indicates a greater abundance of amplifiable cDNA, and presumably mRNA, in the uninfected tick samples versus the infected tick samples.

The difference in *EHMT2* expression, once normalized against the cDNA abundance, shows that infected ticks have 826X higher *EHMT2* expression than uninfected ticks (Table 4). Applying this analysis to individual infected ticks (Figure 4) allows visualization of the variability in *EHMT2* expression. This variability did not correspond in an obvious manner to the (known) suite of tick pathogens other than *B. burgdorferi* or to tick sex (Table 1).

**Figure 4.**
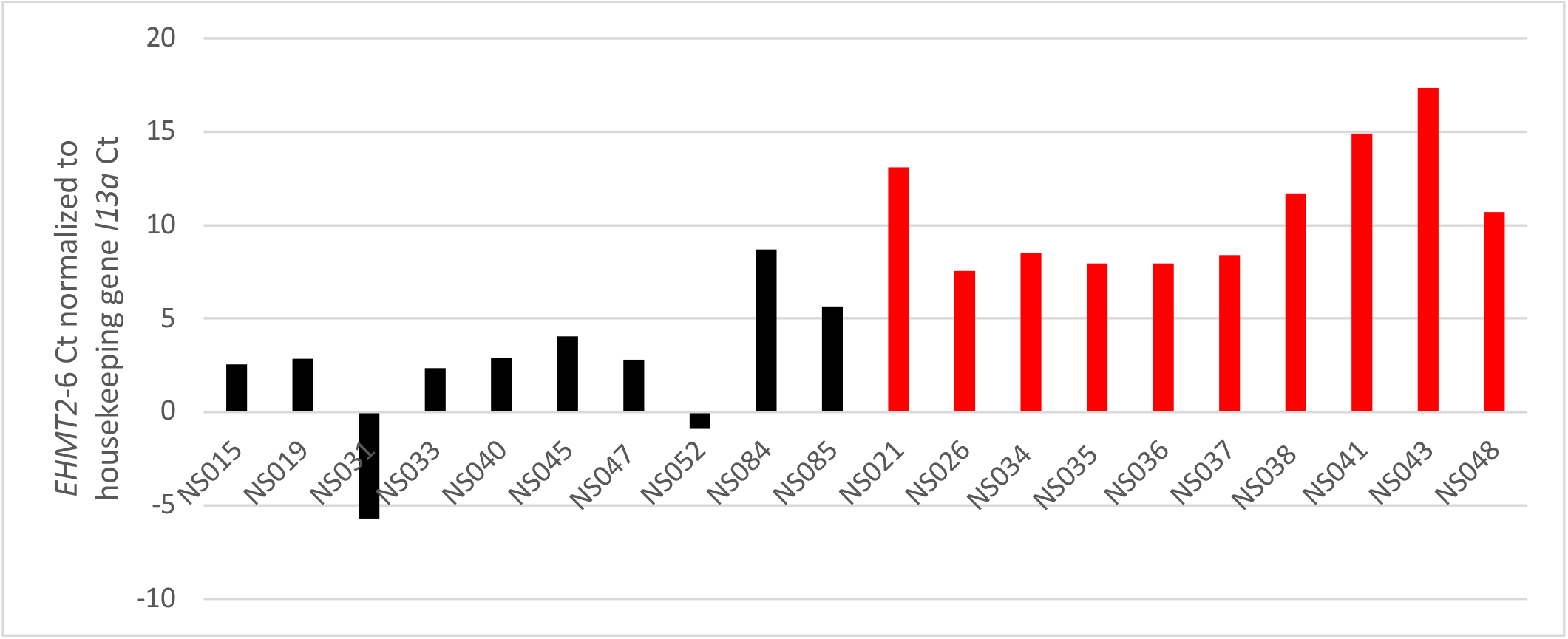
*EHMT2* expression in individual infected ticks (black) relative uninfected ticks (red). Chart shows *EHMT2* expression normalized to the expression of the housekeeping gene *l13a* by subtracting the *l13a Ct* value from the *EHMT2* value; lower differential values are indicative of lower *EHMT2* Ct values representing greater *EHMT2* expression.

## 4.0 Discussion

Epigenetic regulation in ticks has been poorly explored in general, and the role of *B. burgdorferi* in manipulating the epigenome of its *I. scapularis* tick vector remains unexplored [59]. Ticks can vector more diverse pathogens than any other arthropod [23] and in nature ticks frequently transmit multiple pathogens, simultaneously or sequentially [63], [64]. Each of these pathogens interact with each other as well as with the host in a battle to establish infection and find their niche to survive and propagate [63]. Manipulation of the tick vectors via modulation of histone modification enzymes have been documented for *Anaplasma phagocytophilum* [49]. Similarly, tick-vectored Apicomplexa parasites are successful in altering host metabolic activity and signaling pathways [65]. Among these strategies are multiple epigenetic processes: the activation of enzymes regulating epigenetic processes, secretion of proteins that affect chromatin regulatory complexes to repress host genes, and the production of regulatory microRNAs [65]. *B. burgdorferi* has been shown to influence its vector’s transcriptome, physiology and behaviour [37], [66], all of which has implications for human and animal medicine. Despite this, the epigenetic cross talk between the tick vector and this bacterial species remains largely unexplored.

The purpose of this study was to explore the epigenetic effects of *B. burgdorferi* on both DNA modification and histone modifications. *Ixodes scapularis* has DNA methyltransferase enzymes [49] and extensive repeat regions including non-coding pericentromeric tandem repeats of DNA, termed *Ixodes scapularis repeats, ISRs* [23]. *ISR*s have been shown to be methylated based both on differential restriction with methylation-sensitive and methylation-insensitive restriction enzymes [56] and by binding of antibodies against 5-methylcytosine (5mC) to the DNA of the closely related *I. ricinus* tick [55]. We confirmed the presence of 5mC DNA in *I. scapularis* using the sensitive approach of immunoprecipitation of methylated DNA. The detecting of DNA methylation using this approach, in addition to those previously performed [55], [56], adds to the robustness of the conclusion that there is DNA methylation in *I. scapularis*. Further, we show that different *ISR* regions are differentially methylated; *ISR2A, ISR2C*, and *ISR3* are more heavily methylated than *ISR2B* and *ISR2D*. There was, however, no obvious or statistically significant impact of *B. burgdorferi* on DNA methylation of either high, moderate or low methylation *ISRs*. Although not previously detected, low level DNA methylation could be present in coding regions, and our results do not exclude modification of such possible low abundance DNA methylation by the tick microbiome. In contrast, *B. burgdorferi* had a pronounced effect on the transcription of *EHMT2*, a histone modifying “master epigenetic” regulator. In other species this gene has been shown to mediate the organism’s response to physiological stress resulting from infection [1], [13], [14], [15], [16], [17]. Our results showed that ticks infected with *B. burgdorferi* had significantly higher *EHMT2* transcription than uninfected individuals. Interestingly, there was considerable individual variation between ticks.

In many species, DNA methylation is crucial for the stability of the genome, a process encompassing DNA repair and replication, as well as modulating gene expression [67]. However, DNA methylation is variably used in different species and DNA methylation has been found to be generally low across arthropod species [68], [69], [70].

DNA methylation patterns in response to infection by bacteria have been investigated in insects, and to a lesser extent, in arachnids. In some insects, bacterial pathogens have been found to alter the expression of the DNA methylation machinery, promoting bacterial persistence and replication [71], [72], [73], [74], [75]. Similarly, histone modifications and the activity of histone modifying enzymes have been found to be altered in response to bacterial infection [12], [76], [77]. These findings supports a role for epigenetics in regulating the interaction between host or vector and pathogen [78]. Research on the role of DNA methylation in pathogen response in arachnids is more limited than in insects. Abudukadier *et al*., [79] found that linyphiid spiders, *Hylyphantes graminicola*, infected with *Wolbachia* had decreased DNA methylation relative to uninfected spiders, suggesting a potential relationship between bacterial infection and DNA methylation in at least one arachnid. As *I. scapularis* ticks seem to primarily methylate non-coding centromeric repeats [55], [56], in this species, a direct role of DNA methylation in controlling the transcription of genes involved in the immune response is unlikely. However, in *Arabidopsis thaliana* centromeric heterochromatin, including pericentromeric tandem repeats, becomes hypomethylated in response to infection by *Pseudomonas syringae* resulting in cell death and morphological abnormalities [80], defects attributed to disrupted transcriptional activity as genome-wide methylation dynamics [81]. Further, the repeats might control methylation-responsive transcription of non-coding RNAs with regulatory roles. DNA methylation is also fundamentally involved in chromosome stability [67]. In this study, we documented differential DNA methylation between the different *I. scapularis* pericentromeric repeats. While the function of the *ISR* family of non-coding peri-centromeric tandem repeats remains unknown [55], [56], a role in chromosome stability and segregation is plausible [82], which might implicate DNA methylation as having a structural role in the genome. Nevertheless, regardless of the biological role of DNA methylation in the *ISR* repeats, this methylation was not influenced by the presence of *B. burgdorferi*. This led to an expansion of the study into an investigation of differential histone modification in response to pathogens.

### 4.2 EHMT2 Expression in Ticks

To investigate an epigenetic effect of *B. burgdorferi* infection upon histone modifications in the tick *I. scapularis* we examined the transcription of the *EHMT2* epigenetic regulator gene. In the absence of prior knowledge of specific histone modifications and gene targets, although an indirect approach, this was the most feasible approach to this question. Microbes such as *Borrelia, Babesia* and *Anaplasma*, all of which were present in at least some of the ticks used in this experiment (Table 1), have been shown to increase tick fitness [83], [84]. *EHMT2* expression is postulated to modulate stress response to promote viability and/or pathogen tolerance [1], [13], [15], [16], [17]. Based on the known role of *EHMT2* in tolerance and the coevolutionary relationship between ticks and the microbes they vector, we had hypothesized that there would be either no effect on *EHMT2* expression if the pathogens did not provoke any physiological stress in the tick vector, or if there was an effect, *EHMT2* expression would be higher in infected ticks than uninfected ticks. Indeed, we found that infected ticks have *EHMT2* expression levels an average of 826 times higher than uninfected ticks.

The Jak-Stat (Janus Kinase-Signal transducers and activators of transcription) pathway is an evolutionarily conserved signalling pathway that is central to cell communication and function, including homeostatic processes like immunity [85]; deficiencies of the pathway are related to increased susceptibility to infection and decreased immune function [86]. Merkling and colleagues [13] found that *EHMT2* regulated *Drosophila* responses to viral infections via the Jak-Stat pathway and that *EHMT2* deficient mutants had decreased survival after infection. This implies that increased *EHMT2* is protective. Ticks, too, have a functional Jak-Stat signalling pathway that functions in antibacterial responses [87]. That we found that *Borrelia*-infected ticks had significantly greater *EHMT2* expression than uninfected ticks implies that the entire stress response and pathogen tolerance pathway may be conserved between insects and arachnids.

The most obvious limitation of this study is that while we detected a pronounced difference in the expression of *EHMT2* in *B. burgdorferi* – infected versus uninfected ticks, we did not confirm that this change in the transcription of a histone modifying enzyme results in differential histone modifications. This work will require a list of differentially expressed genes in *B. burgdorferi* – infected vs uninfected ticks. Another consideration is that the ticks used for this study were obtained from the natural environment, either after feeding from a host, or wild-caught while questing. As is true of wild populations, these ticks carried a variety of pathogens, some of which may also alter gene expression. The rewiring of tick metabolic networks by *Anaplasma phagocytophilum* has been compellingly documented [88]. Additionally, these ticks were exposed to different, and largely unknown, environmental influences, which could have influenced expression of *EHMT2*. Indeed, the magnitude alterations in *EHMT2* transcription was highly variable between ticks. While using lab-reared ticks with controlled infections could reduce variability, such results would be less applicable to ticks in nature, the more interesting and important question. As epigenetic modifications are inter-related [3], [4], [11], [12], future work will need to examining non-coding RNAs that may be differentially expressed under these conditions. The immune response is altered in ticks infected with *B. burgdorferi* [18], however, the mechanisms involved in mediating the organismal response to pathogens, and the role of epigenetics, remains to be explored.

### 4.4 Significance and Conclusions

Infection by pathogenic *Borrelia* bacteria is an increasing concern for human and animal health. Investigation of the effect of these spirochete bacteria on their mammalian hosts are increasing, nevertheless, this understanding needs to be coupled to an understanding of the interaction between the tick vector and the bacteria to understand the epigenetic negotiations between pathogen and vector. An influence of *Borrelia* bacteria on the physiology and behavior of *Ixodes scapularis* ticks has been documented [39], [44], [83], [84]. The missing link, however, is the mechanisms that *Borrelia* uses to modulate these changes in vector physiology and behavior. Given the pivotal role of epigenetics in mediating environmental influences on the genome, it seemed biologically important to investigate the epigenetic effects of *Borrelia burgdorferi* on *Ixodes scapularis*.

We found that the pericentric repeats of *Ixodes scapularis* are methylated, as previously reported. In addition to confirming this finding, we detected different levels of DNA methylation in individual repeats. The level of DNA methylation, was not, however, significantly responsive to the presence or absence of *B. burgdorferi* in the ticks. In contrast, our results clearly demonstrate that *EHMT2* is significantly over-expressed in infected ticks. To our knowledge, this study is the first to document epigenetic effects of *B. burgdorferi* on *I. scapularis*. This will open an avenue to further understanding of the epigenetic mechanisms through which ticks maintain and transmit pathogens. Further, epigenetic changes can be modified by chemical means, providing a potential avenue to decrease the risk of tick-vectored infections.

## Methods

### Experimental approach

To determine *Borrelia burgdorferi* presence in *Ixodes scapularis* ticks, we assessed differences in both DNA methylation status and expression of *EHMT2/G9a* in *B. burgdorferi-*infected and uninfected ticks. Ticks were obtained as donations from the public and tested for *B. burgdorferi* infection by nested PCR upon receipt. DNA from these ticks were then rescreened for other common tick-borne pathogens. The methylation status of *ISR* repeats, previously determined to be methylated [55], [56], was assessed by magnetic-methylated-DNA immunoprecipitation (MagMeDip) reactions. Expression of *EHMT2/G9a* in *B. burgdorferi-* infected and uninfected ticks was determined by real-time reverse transcription PCR using primers designed for *I. scapularis G9a* and a housekeeping gene.

### Tick Samples

Ticks used for methylation analysis were sourced from archived samples at the Mount Allison Tick Bank (Mount Allison’s Animal Care Committee, #102550). These ticks had been donated by members of the public. Upon receipt, half of each tick was used for DNA testing for tick-borne pathogens as described by Wills et al., (2018) and the other half archived at -20°C. For this study, 26 engorged adult female archived half-ticks, 6 donated in 2018 as part of a pilot study and 20 donated in 2020 were selected (Table 1). Ticks used for the *EHMT2/G9a* expression analysis needed to be collected alive to preserve RNA integrity. These ticks were collected in October 2022, from the Annapolis Valley, Nova Scotia, by collaborator Glen Parsons. Ticks were collected by flagging, placed in a urine specimen container with a moist towel and sent by personal courier to the lab. In the lab, ticks were stored in a 10°C incubator for no more than 7 days. Living ticks were manually killed by dismembering and half ticks were placed in RNALater (Sigma) and stored at -80°C to reduce RNA degradation. Both female and male ticks were used.

Screening of tick samples for tick-borne pathogens: Screening for pathogens common in the region, including *Borrelia*, was conducted. DNA was extracted from the archived half-tick sample or the half tick that was not used for RNA isolation. DNA extraction was as described by [89] with the exception that the DNeasy® Blood & Tissue Kit (Qiagen) was used, following the manufacturer’s recommended protocol for tissue extraction with incubation of the tick samples overnight at 56°C with proteinase K. The concentration of the DNA samples was determined using a Nanodrop®ND-1000 UV-Vis Spectrophotometer (Thermo Fisher Scientific). Tick DNA samples used for methylation testing were selected based on high DNA concentration, a 260/280 ratio between 1.8 and 2.0 and the 260/230 ratio greater than or equal to 2.0.

Nested Polymerase Chain Reaction (nPCR) was performed to detect DNA from *B. burgdorferi, B. miyamotoi, Anaplasma spp*. and *Babesia spp* in the tick DNA. The primers used for pathogen detection and annealing temperatures are shown in Table 2 and 25μl reactions using GoTaq Green (Promega) were performed. All reactions were done in UV-treated dead air PCR cabinets and DNA extraction, PCR and gel electrophoresis were done in separate areas with unconnected air flow. No template controls (using 2μL of nuclease-free water (nfH_2_O) instead of template DNA) were done at the start and end of all PCR manipulations. Amplification involved an initial denaturation step for four minutes at 94°C, denaturation occurred at 94°C for one minute, annealing at the temperatures shown in Table 2 for one minute, and extension at 72°C for one minute, for 35 cycles. The final elongation step was at 72°C for ten minutes. Samples were held at 4°C until removal from the thermocycler and stored at -20°C until further use. The second round of the nPCR was performed similarly using 2μL of the first-round PCR samples as DNA input. Amplicons were visualized using 1.2% agarose gel electrophoresis in sodium borate (SB) buffer with Eco-stain (Bio Basic), imaged using a UV transilluminator (Labnet).

### Primer design

ISR primer design: Primers were designed to amplify 3 families of heterochromatic tandem repeats, *Ixodes scapularis* repeats (*ISR): ISR1, ISR2A, ISR2B, ISR2C, ISR2D*, and *ISR3* previously identified as being methylated [56]. The full nucleotide sequence of each repeat (Table 3) was identified in the NCBI GenBank from the consensus sequences reported in the [56] study, using the basic local alignment search tool (nBLAST). Because of the complexity of designing primers for tandem repeats with several internal repeat motifs, each of the 7-target genomic region primers were manually designed with suitable length (18 to 24 nucleotides), GC content of 50 ± 4%, melting temperature (Tm) of 60°C ± 3°C and a target amplicon length of 200-300 bp, as required for the MeDIP reactions. Primer sets were assessed using an Oligonucleotide properties calculator to screen for hairpins, self-complementarity, and self-annealing sites [90] and in silico PCR using Amplify4. As flaws in the primers were identified, this process was reiterated. Primer sets were then tested with tick DNA and produced amplicons of the expected sizes, albeit with laddering and smearing as would be expected for repeat DNA (Supplemental Figure 1).

Primer design for *I. scapularis EHMT2/G9a* and reference gene: Several primers were designed for this study. Merkling et al. [13] had previously published the sequence *of Drosophila melanogaster EHMT2*, which was used to search *Ixodes scapulari*s genome sequences to find the most similar *Ixodes scapulari*s gene (Accession number: GHJT01005725), which was presumed to be *Ixodes scapulari*s *EHMT2*. The NIH Primer BLAST tool was used to generate primers for the *Ixodes scapularis* predicted *EHMT2* mRNA, of which, two were chosen (Table 2).

### PCR optimization

*ISR* primers: The optimal annealing temperature for each primer pair was determined empirically through conventional end point gradient PCR using GoTaq Green (Promega). Gradient temperatures tested for ISR1 were: 51, 51.9, 53.4, 55.6, 58.7, 60.9, 62.3, 63, 63.4, 65, 67.2, 68 °C. Gradient temperatures tested for ISR2A, ISR2B, and ISR2C were: 47.4, 48.6, 50.4, 51, 51.9, 53.4, 55.6, 58.7, 60.9, 62.3, 63, 63.4, 65, 67.2, 68 °C. ISR3 gradient temperatures were: 51, 51.9, 53.4, 55.6, 58.7, 60.9, 62.3, 63 °C. Amplification conditions were: a 3-minute denaturing period at 95°C, followed by 40 cycles of 30 second denaturation at 95°C, a 60-seconding annealing period and a 30 second elongation period at 72°C followed by a final elongation step at 72°C for 10 minutes. Amplification results were assessed by agarose gel electrophoresis as described above and the optimal amplification temperature is shown in Table 2. While a single clear amplicon is optimal, repeat sequences can generate ladders or smears upon amplification. The predicted amplicon size of ISR1 is 215 bp, and a faint band at this size is seen (Supplemental Figure 1). During subsequent qPCR reactions, however, amplification was found to be superior at 51°C, so this annealing temperature was used. An empirically determined annealing temperature for ISR 2A and 2B primers was not obtained; both consistently produced a smear (Supplemental Figure 1) so the predicted annealing temperature of 55°C was used for qPCR. The predicted amplicon sizes of ISR2C and ISR2D are 242 and 213 bp, respectively; a faint band consistent with these sizes, embedded in a smear of different length fragments, was seen (Supplemental Figure 1). ISR3 produced multiple bands, the most prominent of which was consistent with the expected 211 bp amplicon size (Supplemental Figure 1). This band remained prominent at 55°C so this temperature was used for convenience.

Optimization of *EHMT2* primers: Koči et al. [91] documented that *Ixodes scapularis rps4* and *l13a* were effective reference genes with constant expression through development and external environmental perturbations, so primers for these genes (Table 2) were designed as described above for normalization of *EHMT2* gene expression. Amplification of products consistent with the expected products are shown in Supplemental Figure 2. The *l13a* primer set was the most sensitive (lowest Ct values) so that primer set was used for this study (Supplemental Table 1). Amplification conditions for the control genes were described by [91] and since the annealing temperature of both designed *EHMT2* primers were only 5°C higher than the annealing temperatures of the reference genes, this PCR program was used for the *EHMT2* primers as well. If gel electrophoresis did not immediately follow qPCR, qPCR products were stored at -20°C. Amplification conditions consisted of 95°C for 2 minutes followed by 40 cycles of denaturation at 95°C for 15 seconds, annealing at 55°C for 30 seconds and elongation at 72°C for 30 seconds.

qPCR calibration, qPCR conditions and standard curves: The iCycler, Bio-Rad was calibrated prior to measurements according to the manufacturer’s instructions with FAM dye and external well factor solution (Bio-Rad). All qPCR preparations were performed in a PCR hood (Fisher Scientific PCR Workstation) using autoclaved and UV-irradiated equipment and consumables. qPCR reactions were prepared in triplicate or duplicate for the *ISR* amplifications and in duplicate for the *EHMT* study. Reactions were prepared in 8-strip PCR tubes or 96 well plates (BrandTech, Cat. 781320 or 781375), with 10 μL of iQTM SYBR Green Supermix (BioRad, Cat. 1708880), 1 μL forward primer and 1 μL reverse primer specific to the region being targeted, 10 μL nf-H2O and 2 μL of input DNA for the *ISR* amplifications, and half as much (10μl total reactions) for the *EHMT* samples (previously validated to perform as well as the 20μl reactions). Two “no template” negative controls lacking input DNA were included in each run for each primer set.Following calibration, standard curves were produced to allow correlation of Ct values for each primer set to known amounts of input DNA. For the *ISR* primers, 2-fold serial dilutions of tick DNA from 8ng/μl to 0.125ng/μl using IQ™ SYBR® Green Supermix (Bio-Rad, C#64313832) were used. For *EHMT2* and *l13a* primers a series of standard dilutions with both infected and uninfected tick cDNA was performed. Only the lowest three dilutions (1:1, 1:2, 1:4) generated Ct values (Supplemental Figure 3). Standard curves, plotting the Ct values against DNA concentrations, were made for each *ISR* region, with the exception of *ISR1* which failed to amplify reliably at 51°C (Supplemental Figure 1). The quality of the standard curve was determined by its R^2^ value, which were all above 0.9 for the *ISR* regions. The standard curve was used to determine the efficiency of each primer, the linear range, and the detection and quantification limit. The slope of the standard curves allows determination of the PCR efficiency by using the formula E=10^−1/slope^. Efficiency of the ISR regions ranged from 2.044 to 1.536; an efficiency value of 2 is optimal (Ruijter et al., 2021). Amplification efficiencies for each of the *ISR* regions, *ISR2A, ISR2B, ISR2C, ISR2D* and *ISR3* were 1.726, 1.536, 2.044, 1.706, and 1.847, respectively. Amplification efficiency was calculated for each primer, yielding 1.88 for *EHMT2 6* and 1.92 for *l13a*. These efficacies satisfy the assumption that there is near equal primer efficiency between both genes and are within 10% of the optimal amplification value of 2.

Determining DNA methylation of *I. scapularis* centromeric repeats by MeDIP: The Diagenode MagMeDip qPCR kit (Diagenode, C02010021) was used to quantify the DNA methylation status of *ISR* centromeric repeats of ticks infected and uninfected with *B. burgdorferi*. This protocol involved a series of 6 steps: (1) Cell collection and lysis; (2) DNA extraction and purification; (3) DNA shearing; (4) Methylated DNA immunoprecipitation; (5); Methylated DNA isolation; (6) qPCR analysis.

The manufacturer’s protocol was followed with the following parameters and exceptions. As the starting material was archived DNA, steps 1 and 2 were not needed. To determine DNA fragment length, the integrity of the archived DNA was first assessed by agarose gel electrophoresis. Samples with initial fragment sizes larger than 400 bp were diluted to 1.2 μg of DNA in 55 μL with the GenDNA TE buffer and sonicated using a sonication bath (FS30, Fisher Scientific, Serial#RTA050266358) set to 8-10°C and sheared with 10 repetitions of 30 second pulses with return to ice for 30 seconds between pulses. Gel electrophoresis was then used to assess fragment size and sonication repeated until fragments of approximately 400bp were produced. The methylated-DNA immunoprecipitations reactions were prepared as described by the manufacturer. IP tubes were placed on a rotating wheel (SCILOGEX SCI-RD-E Analog Tube Rotator Mixer, C#824230019999), set at 40 rpm, at 4°C for approximately 18 hours. The kit-provided meDNA spike-in and unDNA spike-in controls, sheared methylated and unmethylated DNA from *Arabidopsis thaliana*, were used. Aliquots of DNA (1/10^th^ of the IP sample) prepared in the same manner and at the same time as the experimental samples but not subjected to immunoprecipitation, were used for the input samples. Following immunoprecipitation, the controls and immunoprecipitation reactions were again treated in parallel. *Borrelia*-infected and non-infected samples were tested in parallel. The isolation of the methylated DNA was performed as described by the manufacturer and the resulting DNA stored at -20°C until qPCR was performed. Prior to processing the experimental samples, control reactions targeting the 2 spike-in controls were prepared to determine the efficiency of each MeDIP reaction. These reactions were performed in duplicate for each spike-in control as described by the manufacturer. qPCR reactions measuring the percentage of DNA methylation for the immunoprecipitated or input control DNA for each of the *ISR* target regions.

Input DNA and IP DNA were diluted 1:2 with nf H2O and used for subsequent qPCR reactions for each of the *ISR* target regions as described above. The Ct values were used to calculate the percentage of recovery of methylated DNA with the equation: % recovery= E^ [(Ct (10% input) – X) -Ct (IP)] x 100. Where E is the amplification efficiency of the primers and “X” is the value to which the amplification efficiency, E, must be raised to equal 10, to compensate for the 1/10^th^ dilution of the input sample. In cases where E has not been determined, this can be approximated by: % recovery= 2^ [Ct (10% input) – 3.32 -Ct (IP sample)] x 100. In cases where the methylation of the target region is being compared to a reference region of the genome, the enrichment status of the sample can be determined; this was not done in this study as there was no relevant reference gene.

### *EHMT2* gene expression

RNA was extracted using the Qiagen RNEasy Mini Kit following the manufacturer’s directions. All work was performed in a designated sterile RNA hood, instruments decontaminated. The RNAlater in which the dissected ticks were stored was manually removed with a pipette. After addition of Qiagen RLT buffer was added, the dissected tick samples were manually homogenized with a micropestle, then centrifuged (Spectrafuge 24D) for 3 minutes at 16.3X*g* to pellet the debris. The supernatant was removed and further processing was conducted according to the manufacturer’s instructions. Each sample was treated for 15 minutes with the supplied DNase solution. Samples were eluted with 30μL of RNase free water which was applied to the column twice. 3μL of each sample was aliquoted for quantification via NanoDrop (NanoDrop 1000), and the remainder was immediately frozen at -80°C. For each sample, a cDNA synthesis reaction was performed, as well as a “no-RT” control to ensure the removal of residual genomic DNA. All reactions were performed in a sterile Clean Prep Hood using NanoDrop 1000 and SandWatch software (New England Biolabs) and 250ng input RNA in a 20ul reaction according to the manufacturer’s instructions. cDNA synthesis occurred in a themocycler (MultiGene OptiMAX Labnet International, Inc), for a 2 min at 25°C primer annealing step, 10 minutes at 55°C for cDNA synthesis and 95°C for 1 minute for heat inactivation. Samples were then frozen at -20°C until use.

### Statistical Analysis

For assessing the significance of DNA methylation in the ISR repeats, between repeats and in response to *Borrelia* infection, the Ct values determined by the qPCR were used for calculations of the percentage of DNA recovered as described in 2.5. R studio was used to conduct an unpaired t-test measuring DNA methylation differences by infection status and an ANOVA was used to assess the difference in DNA methylation between ISR repeats, although the repeats are not independent as the same ticks are used across primer regions.

To assess the results of *EHMT2* gene expression, the Ct values generated from qPCR were normalized to the reference gene amplification via double delta Ct analysis [62]. This analysis requires that there is near equal efficiency of each primer set (within 5%), there is near 100% amplification efficacy of the reference and target genes, and that internal control genes are consistently expressed regardless of treatment [62]; efficiency of the primers sets are described above and developmental stability of expression of the reference genes is shown by Koci et al. [91].

## Acknowledgements

We would like to extend our gratitude to Glen Parsons for collection of live ticks and the many members of the public and veterinarians for donation of feeding ticks. We thank Samantha Bishop and Ebruvibiyo Daniella Ibru for pathogen testing of the ticks and Julie Lewis for invaluable discussion and feedback on this manuscript. Seed funding for the tick bank was provided by the Canadian Lyme Disease Foundation and this work was supported by a Natural Sciences and Engineering Research Council grant to VKL and undergraduate summer studentship awards to GM, AN and JV.

