## Supplemental tables and figures for "Epigenetic responses in *Borrelia*-infected Ixodes *scapularis* ticks: Over-expression of euchromatic histone lysine methyltransferase 2 and no change in DNA methylation"

**Supplemental Table 1. qPCR results for control housekeeping genes *l13a* and *rps4***

| Sample | Pathogen Infection Status | *l13a* Dup 1 Ct | *l13a* Dup 2 Ct | *l13a* Avg. Ct | *rps4* Dup 1 Ct | *rps4* Dup 2 Ct | *rps4* Avg. Ct |
|---|---|---|---|---|---|---|---|
| NS021 | Negative | 20.7 | 20 | 20.35 | 26.5 | 26.7 | 26.1 |
| NS026 | Negative | 23.2 | 23 | 23.1 | 30.1 | 30.2 | 30.15 |
| NS034 | Negative | 23.6 | 22.9 | 23.25 | 30.6 | 30.2 | 30.4 |
| NS035 | Negative | 25.1 | 26.1 | 25.6 | 33.4 | 34.5 | 33.95 |
| NS036 | Negative | 26.7 | 27.2 | 26.95 | 35.4 | 36.6 | 36 |
| NS037 | Negative | 20.7 | 20.7 | 20.7 | 26.7 | 26.9 | 26.8 |
| NS038 | Negative | 23.6 | 23.5 | 23.55 | 28 | 29.9 | 28.95 |
| NS041 | Negative | 21.9 | 22.2 | 22.05 | 27.8 | 27.9 | 27.85 |
| NS043 | Negative | 22.6 | 22.7 | 22.65 | 28.4 | 28.5 | 28.45 |
| NS048 | Negative | 23.3 | 24 | 23.65 | 31.1 | 30.2 | 30.65 |
| Neg. Cont. | N/A | N/A | N/A | N/A | N/A | N/A | N/A |
| No temp Cont. | N/A | N/A | N/A | N/A | N/A | N/A | N/A |

Note: "N/A" means no Ct value was returned, suggesting no amplification. For calculations, this result was assigned a value of "40".

**Supplemental Table 2.** ***Nanodrop results for RNA extracted form infected and uninfected ticks***

| Sample ID | Infection Status | ng/µL | A260 | A280 | 260/280 | 260/230 | Abs. |
|---|---|---|---|---|---|---|---|
| NS015 | Positive | 68.5 | 1.711 | 1.259 | 1.36 | -0.72 | -2.372 |
| NS019 | Positive | 55.1 | 1.378 | 1.026 | 1.34 | -0.77 | -1.787 |

| | | | | | | | |
|---|---|---|---|---|---|---|---|
| NS031 | Positive | 63.9 | 1.598 | 1.149 | 1.391 | -0.77 | -2.071 |
| NS033 | Positive | 25.6 | 0.641 | 0.487 | 1.32 | -0.16 | -3.892 |
| NS040 | Positive | 21.5 | 0.538 | 0.404 | 1.33 | -0.21 | -2.557 |
| NS045 | Positive | 39.7 | 0.992 | 0.709 | 1.40 | -0.35 | -2.862 |
| NS047 | Positive | 44.4 | 1.110 | 0.759 | 1.46 | 0.18 | 6.332 |
| NS052 | Positive | 40.4 | 1.010 | 0.692 | 1.46 | 0.28 | 3.553 |
| NS084 | Positive | 27.9 | 0.689 | 0.470 | 1.49 | 0.27 | 2.627 |
| NS085 | Positive | 19.6 | 0.491 | 0.336 | 1.46 | 0.22 | 2.228 |
| NS021 | Negative | 19.6 | 0.491 | 0.323 | 1.52 | 0.38 | 1.308 |
| NS026 | Negative | 26.9 | 0.672 | 0.434 | 1.55 | 0.14 | 4.864 |
| NS034 | Negative | 37.0 | 0.925 | 0.649 | 1.43 | 0.37 | 2.525 |
| NS035 | Negative | 29.2 | 0.729 | 0.508 | 1.44 | 0.29 | 2.504 |
| NS036 | Negative | 58.3 | 1.457 | 1.062 | 1.37 | 0.51 | 2.871 |
| NS037 | Negative | 25.7 | 0.642 | 0.471 | 1.36 | 0.36 | 1.800 |
| NS038 | Negative | 28.7 | 0.718 | 0.513 | 1.40 | 0.17 | 4.208 |
| NS041 | Negative | 25.7 | 0.642 | 0.488 | 1.32 | -0.18 | -3.662 |
| NS043 | Negative | 32.8 | 0.819 | 0.591 | 1.39 | -0.29 | -2.870 |
| NS048 | Negative | 71.6 | 1.791 | 1.352 | 1.33 | -8.01 | -0.224 |

**Supplementary Table 3. qPCR results for positive and negative ticks with *EHMT2* and *l13a* primer sets**

| Sample | Pathogen Infection Status | *EHMT2* 6 Dup 1 Ct | *EHMT2* 6 Dup 2 Ct | Avg. *EHMT2* 6 Ct | *EHMT2* 8 Dup 1 Ct | *EHMT2* 8 Dup 2 Ct | Avg *EHMT2* 8 Ct | *l13a* Dup 1 Ct | *l13a* Dup 2 Ct | Avg. *l13a* Ct |
|---|---|---|---|---|---|---|---|---|---|---|
| NS015 | Positive | N/A | 37.9 | N/A | N/A | 37.7 | N/A | 35.3 | 37.6 | 36.45 |
| NS019 | Positive | 35.2 | 34.6 | 34.9 | 38.4 | 35 | 36.7 | 31.6 | 32.5 | 32.05 |
| NS031 | Positive | 34.6 | 34 | 34.3 | 33.7 | N/A | N/A | N/A | 30.7 | N/A |
| NS033 | Positive | 31.4 | 31 | 31.2 | N/A | 35.3 | N/A | 28.3 | 29.4 | 28.85 |
| NS040 | Positive | 35.1 | 34.6 | 34.85 | 32.2 | N/A | N/A | 32.2 | 31.7 | 31.95 |
| NS045 | Positive | 34.8 | 33.8 | 34.3 | 36.3 | N/A | N/A | 30.5 | 30 | 30.25 |

| | | | | | | | | | | |
|---|---|---|---|---|---|---|---|---|---|---|
| NS047 | Positive | 34.6 | 34.9 | 34.75 | 36.8 | 34.5 | 35.65 | 31.7 | 32.2 | 31.95 |
| NS052 | Positive | 34.9 | 33.3 | 34.1 | N/A | N/A | N/A | 29.2 | 30.8 | 35 |
| NS084 | Positive | 35.4 | N/A | N/A | 36.8 | 37.5 | 37.15 | 28.8 | 28.8 | 28.8 |
| NS085 | Positive | 33.7 | 31.3 | 32.5 | 32.7 | N/A | N/A | 27.8 | 25.9 | 26.85 |
| Neg. Cont. | N/A | 34.5 | N/A | N/A | N/A | N/A | N/A | N/A | N/A | N/A |
| No temp. Cont. | N/A | N/A | N/A | N/A | N/A | N/A | N/A | N/A | N/A | N/A |
| NS021 | Negative | 35.1 | 31.8 | 33.45 | 36.1 | 35.1 | 35.6 | 20.7 | 20 | 20.35 |
| NS026 | Negative | 30.9 | 30.4 | 30.65 | N/A | 36.4 | N/A | 23.2 | 23 | 23.1 |
| NS034 | Negative | 32.7 | 30.8 | 31.75 | 37.1 | 37.7 | 37.4 | 23.6 | 22.9 | 23.25 |
| NS035 | Negative | 34 | 33.1 | 33.55 | 37.3 | 38.2 | 37.75 | 25.1 | 26.1 | 25.6 |
| NS036 | Negative | 35.7 | 34.1 | 34.9 | N/A | N/A | N/A | 26.7 | 27.2 | 26.95 |
| NS037 | Negative | 29.5 | 28.6 | 29.1 | 36.7 | 36 | 36.35 | 20.7 | 20.7 | 20.7 |
| NS038 | Negative | 32.6 | 37.9 | 35.25 | 37.4 | 35.5 | 36.45 | 23.6 | 23.5 | 23.55 |
| NS041 | Negative | 36.8 | 37.1 | 36.95 | 37.9 | 36.6 | 37.25 | 21.9 | 22.2 | 22.05 |
| NS043 | Negative | N/A | 34.7 | N/A | N/A | 37.9 | N/A | 22.6 | 22.7 | 22.65 |
| NS048 | Negative | 37.4 | 31.3 | 34.35 | 33.3 | 36.5 | 34.9 | 23.3 | 24 | 23.65 |
| Neg. Cont. | N/A | N/A | N/A | N/A | N/A | N/A | N/A | N/A | N/A | N/A |
| No temp Cont. | N/A | N/A | N/A | N/A | N/A | N/A | N/A | N/A | N/A | N/A |

Note: “N/A” means no Ct value was returned, suggesting no amplification. For calculations, this result was assigned a value of “40”.

**Supplemental Table 4. No-RT controls**

| Sample | Infection Status | *EHMT2* 6 Dup 1 Ct | *EHMT2* 6 Dup 2 Ct | | *EHMT2* 8 Dup 1 Ct | *EHMT2* 8 Dup 2 Ct | | *l13a* Dup 1 Ct | *l13a* Dup 2 Ct | Avg. |
|---|---|---|---|---|---|---|---|---|---|---|

| | | | | Avg *EHMT2* 6 Ct | | | Avg. *EHMT2* 8 Ct | | | *l13a* Ct |
|---|---|---|---|---|---|---|---|---|---|---|
| NS015 | Positive | N/A | N/A | N/A | N/A | N/A | N/A | N/A | 37.6 | N/A |
| NS019 | Positive | 33.7 | N/A | N/A | N/A | N/A | N/A | N/A | N/A | N/A |
| NS031 | Positive | 34.5 | 36.1 | 35.31 | N/A | N/A | N/A | N/A | N/A | N/A |
| NS033 | Positive | N/A | N/A | N/A | N/A | N/A | N/A | N/A | N/A | N/A |
| NS040 | Positive | N/A | N/A | N/A | N/A | N/A | N/A | N/A | N/A | N/A |
| NS045 | Positive | 38 | 37.2 | 37.6 | N/A | 35.3 | N/A | N/A | 37.6 | N/A |
| NS047 | Positive | N/A | N/A | N/A | N/A | N/A | N/A | 35 | 35.1 | 35 |
| NS052 | Positive | N/A | N/A | N/A | N/A | 35.4 | N/A | N/A | 35.4 | N/A |
| NS084 | Positive | N/A | N/A | N/A | N/A | N/A | N/A | N/A | N/A | N/A |
| NS085 | Positive | N/A | N/A | N/A | N/A | N/A | N/A | N/A | N/A | N/A |
| Neg. Cont. | N/A | N/A | N/A | N/A | N/A | N/A | N/A | N/A | N/A | N/A |
| No temp. Cont | N/A | N/A | N/A | N/A | N/A | N/A | N/A | N/A | N/A | N/A |
| NS021 | Negative | 38.5 | N/A | N/A | N/A | N/A | N/A | N/A | N/A | N/A |
| NS026 | Negative | N/A | N/A | N/A | N/A | N/A | N/A | N/A | N/A | N/A |
| NS034 | Negative | N/A | N/A | N/A | N/A | N/A | N/A | N/A | N/A | N/A |
| NS035 | Negative | N/A | N/A | N/A | N/A | N/A | N/A | N/A | N/A | N/A |
| NS036 | Negative | N/A | N/A | N/A | N/A | N/A | N/A | N/A | N/A | N/A |
| NS037 | Negative | N/A | N/A | N/A | N/A | N/A | N/A | N/A | N/A | N/A |
| NS038 | Negative | 31.4 | 31 | 31.2 | 35.8 | 35.8 | 35.8 | 28.3 | 28.9 | 28.6 |
| NS041 | Negative | N/A | N/A | N/A | N/A | N/A | N/A | N/A | N/A | N/A |
| NS043 | Negative | N/A | N/A | N/A | N/A | N/A | N/A | N/A | N/A | N/A |
| NS048 | Negative | N/A | N/A | N/A | N/A | N/A | N/A | N/A | N/A | N/A |

| | | | | | | | | | | |
|---|---|---|---|---|---|---|---|---|---|---|
| Neg. Cont. | N/A | N/A | N/A | N/A | N/A | N/A | N/A | N/A | N/A | N/A |
| No temp. Cont. | N/A | N/A | N/A | N/A | N/A | N/A | N/A | N/A | N/A | N/A |

Notes: N/A indicates that no Ct vale was obtained, suggesting no amplification occurred. qPCR results indicated some anomalies in qPCR readings, but these were not consistent across primers and no banding was present on gels after gel electrophoresis (Sup Figure 6) with *EHMT2 6* and *l13a* primers, so all samples were included in the study.

Supplemental Figure 1

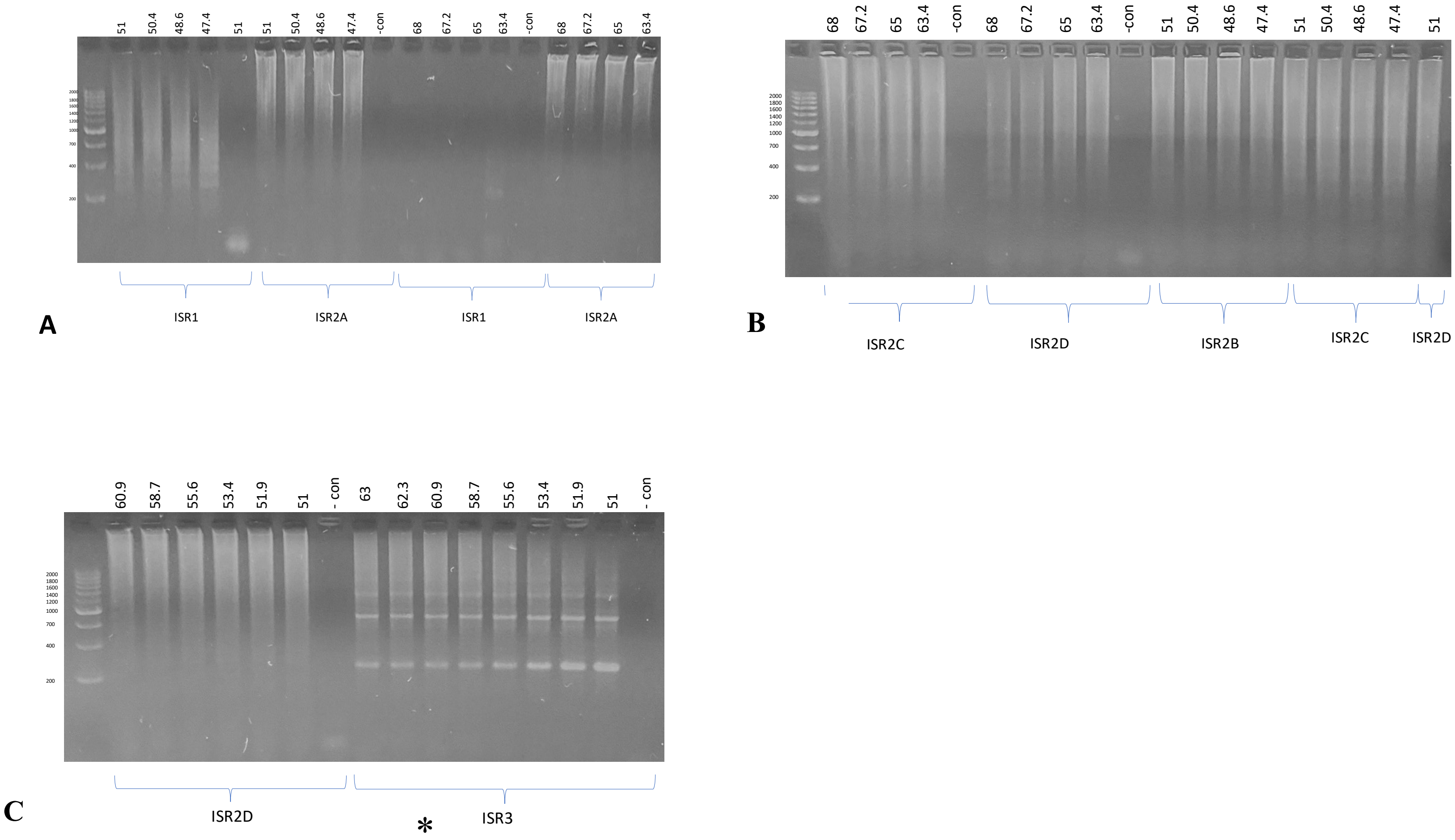


**Supplemental Figure 1:** Agarose gel of Gradient PCR products for ISR primers. All primer optimizations steps were tested on tick DNA samples #590 and #592 from 2019. "– con" = no template negative control using water instead of DNA template. The 200-2000 base pair DNA ladder is to the left in each gel image. A: Gradient PCR products targeting the ISR1 region. B: Gradient PCR products targeting the ISR2C and ISR2D regions. C: Gradient PCR products targeting the ISR3 region. While primers ideally produce a single amplicon, as these are repeat regions primers ISR2C, ISR2D, and ISR3 produced multiple bands.

Supplemental Figure 2

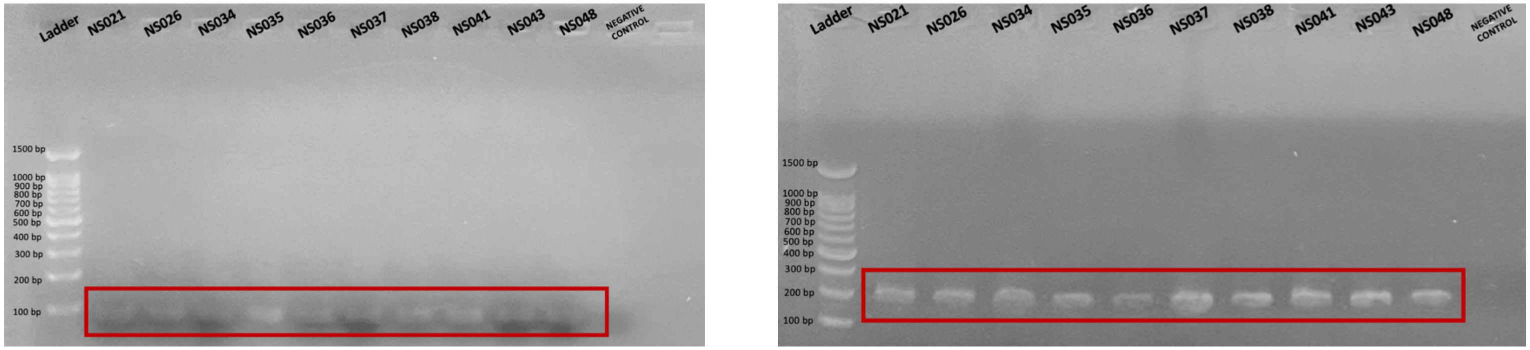


**A** **B**

Supplemental Figurue 2: Gel electrophoresis of qPCR amplification of cDNA from samples NS021, 026, 034, 035, 037, 038, 041, 043, 048 negative ticks. A) *rps4* primers, with an amplicon consistent with the predicted size of 80 bp B) *l13a* primers, with an amplicon consistent with the predicted size of 280 bp.

Supplemental Figure 3

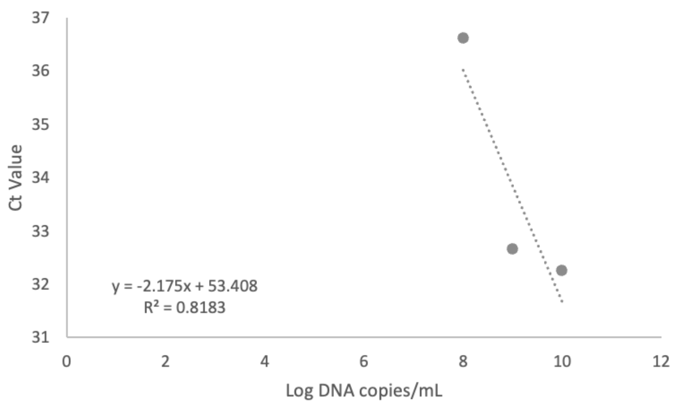


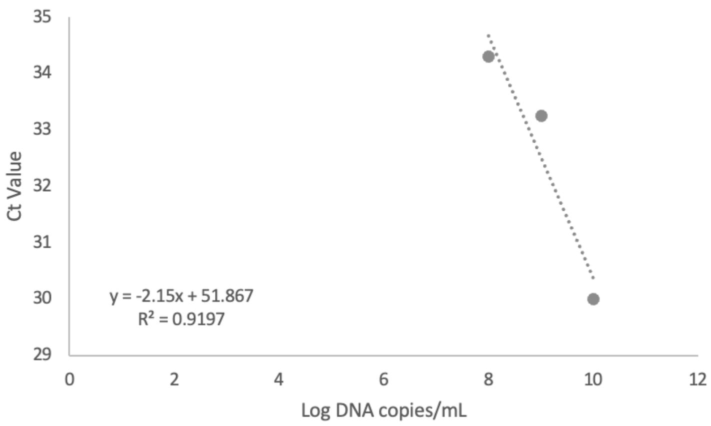


| Dilution | Avg *EHMT2* 6 Ct | Avg *l13a* Ct |
|---|---|---|
| 1 | 32.25 | 30 |
| 0.5 | 32.65 | 33.25 |
| 0.25 | 36.6 | 34.3 |
| 0.125 | 36.65 | 36.25 |
| 0.0625 | 37.25 | 38.65 |
| 0.0312 | 38.7 | N/A |
| 0.016 | N/A | N/A |
| 0.0078125 | 36.15 | 38.95 |
| 0.00390625 | 38.85 | N/A |
| 0 | 37.25 | N/A |

EHMT2 6 *and* l13a *standard curves.* A) Scatterplot with trendline for a three series dilution of *EHMT2 6* primers. The slope of the trendline was calculated to be -2.175, with an $R^2$ value of 0.8183. The amplification efficiency was thus calculated to be 1.88, or 188%. B) Scatterplot with trendline for a three series dilution of *l13a*. The slope of the trendline was calculated to be -2.15, with an $R^2$ value of 0.9197. The amplification efficiency was thus calculated to be 1.92, or 192%. C) qPCR results

Supplemental Figure 4

**A**

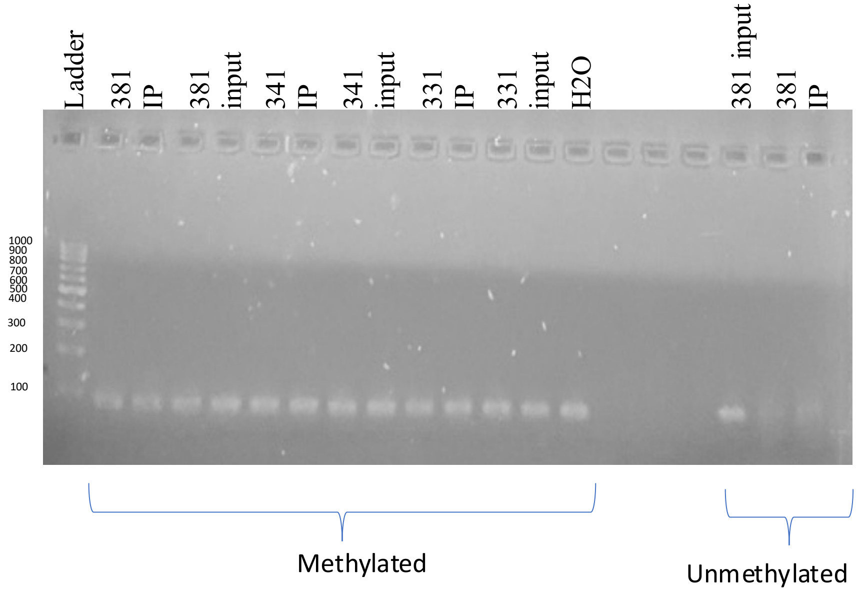

**B**

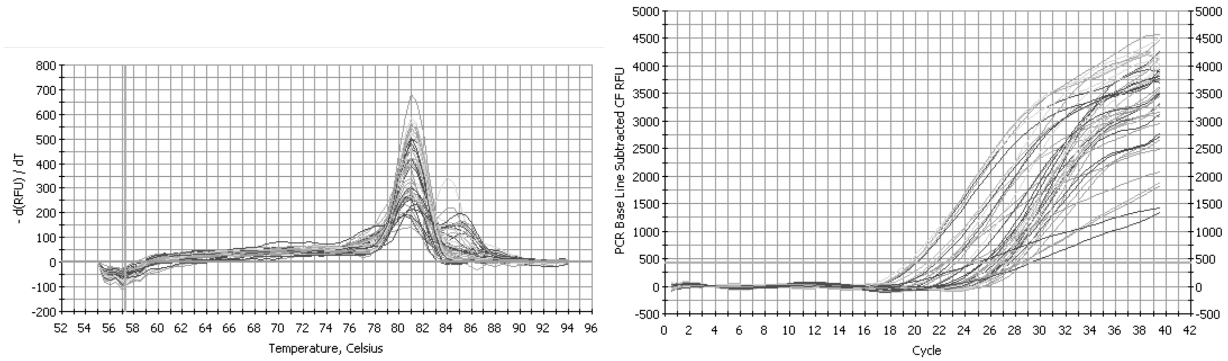


**C**

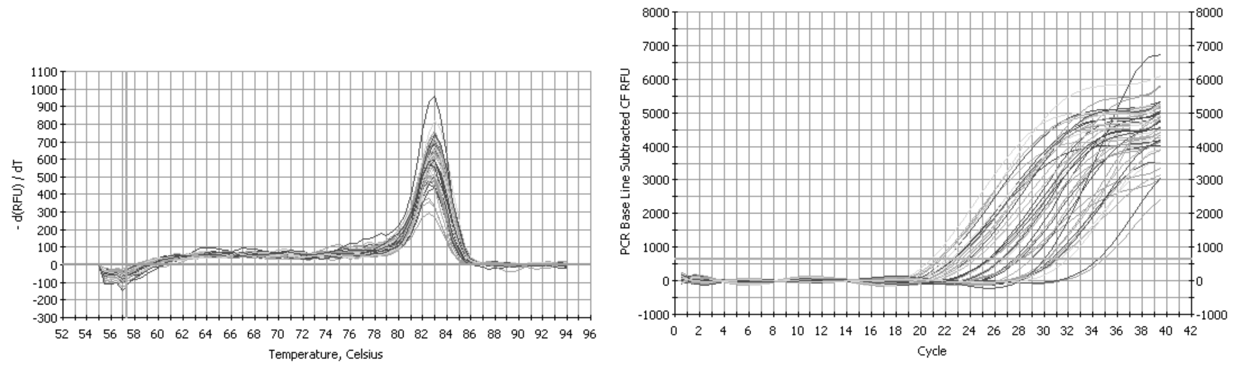


**Supplemental Figure 4:** Control qPCR reaction to determine MeDIP efficiency using the spike-in control DNA and primers. The 200-2000 bp ladder was used for A as a DNA size reference. A: Methylated and unmethylated spike-in control qPCR gel results. Water was used as the negative control. B: qPCR melt curve and amplification curve for the unmethylated spike-in control. C: qPCR melt curve and amplification curve for the methylated spike-in control. The well labels on the gel represent the identity of the tick with IP being immunoprecipitation and the input DNA from the tick samples. The methylated and unmethylated spike-in DNA were used corresponding with the methylated and unmethylated primers.

Supplemental Figure 5

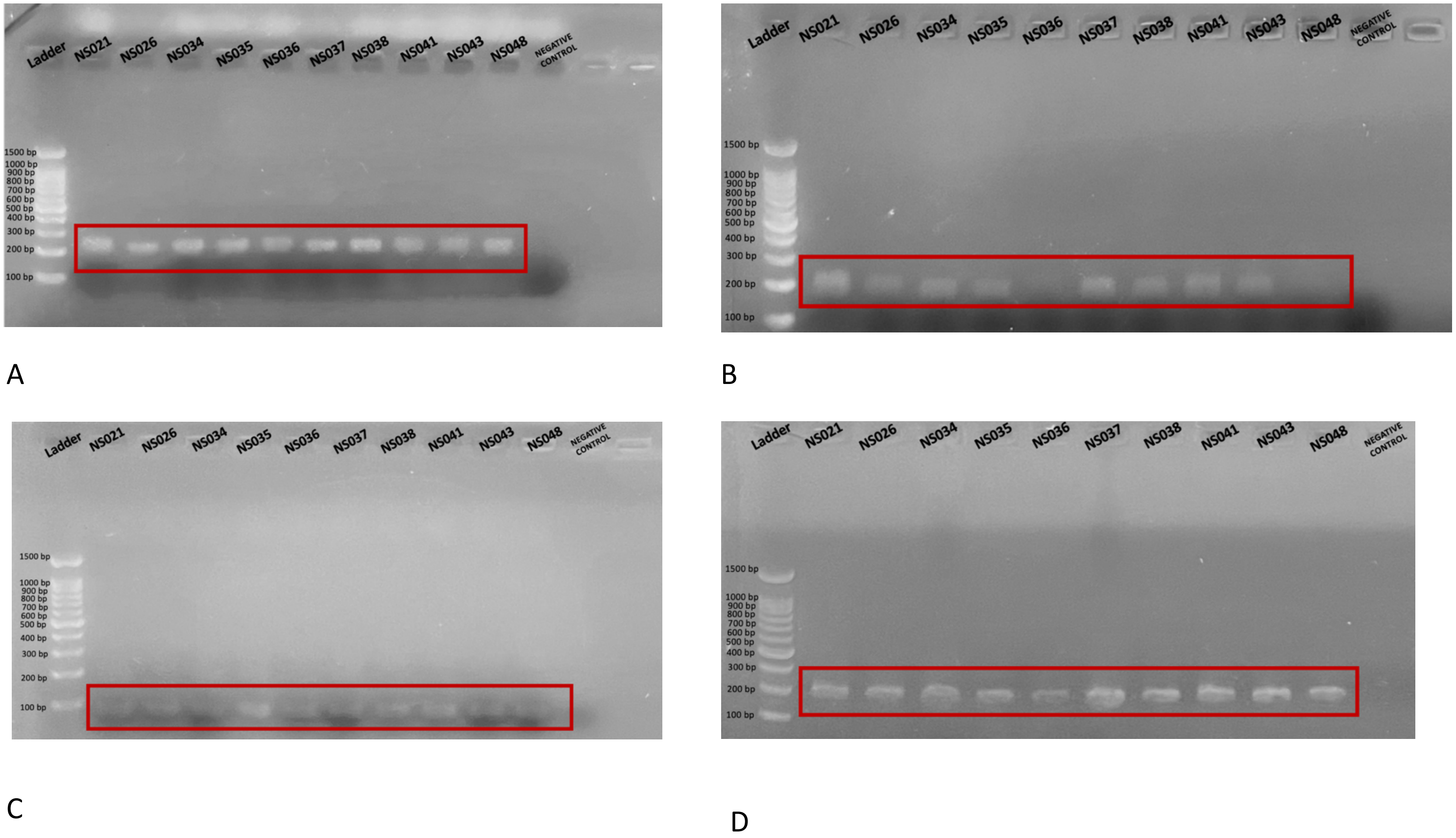


Supplemental Figure 5: Agarose gel electrophoresis of qPCR results A) Synthesized cDNA from each sample underwent qPCR with *EHMT2* 6 primers, with an amplicon size of 207bp. B) Synthesized cDNA from each sample underwent qPCR with *EHMT2* 8 primers, with an amplicon size of 184bp. C) Synthesized cDNA from each sample underwent qPCR with *rps4* primers, with an amplicon size of 80 bp. D) Synthesized cDNA from each sample underwent qPCR with *l13a* primers, with an amplicon size of 280 bp.

Supplemental Figure 6

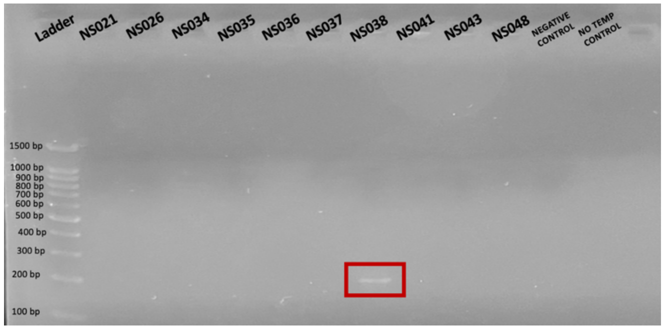


A

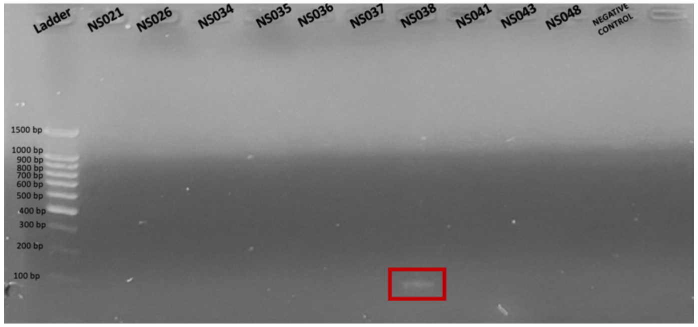


B

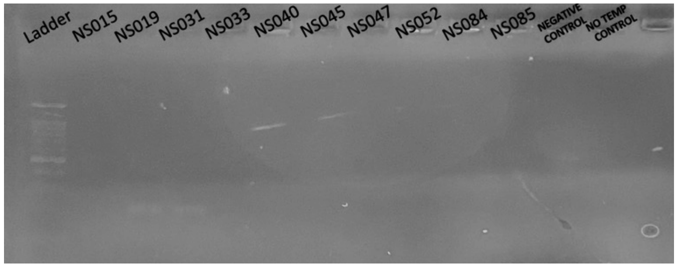


C

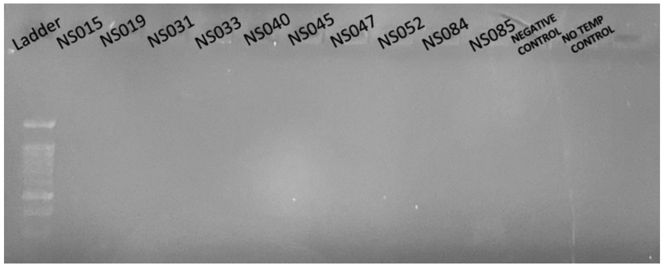


D

Supplemental Figure 6: Agarose gel electrophoresis of qPCR products from no-RT control samples. A) No-RT controls from cDNA synthesis for negative samples that underwent qPCR with *EHMT2* 6 primers, with an amplicon size of 207bp. One sample, NS038 was found to contain residual genomic DNA (box), whereas the rest contained no genomic DNA. B) No-RT controls from cDNA synthesis for negative samples that underwent qPCR with EHMT2 8 primers, with an amplicon size of 184bp. NS038 was again found to contain residual genomic DNA (box), whereas the rest contained no genomic DNA. C) No-RT controls from cDNA synthesis for each positive tick sample underwent qPCR with *EHMT2* 6 primers, with an amplicon size of 207bp. D) No-RT controls from cDNA synthesis for each positive tick sample underwent qPCR with *l13a* primers, with an amplicon size of 280bp.

Supplemental Figure 7

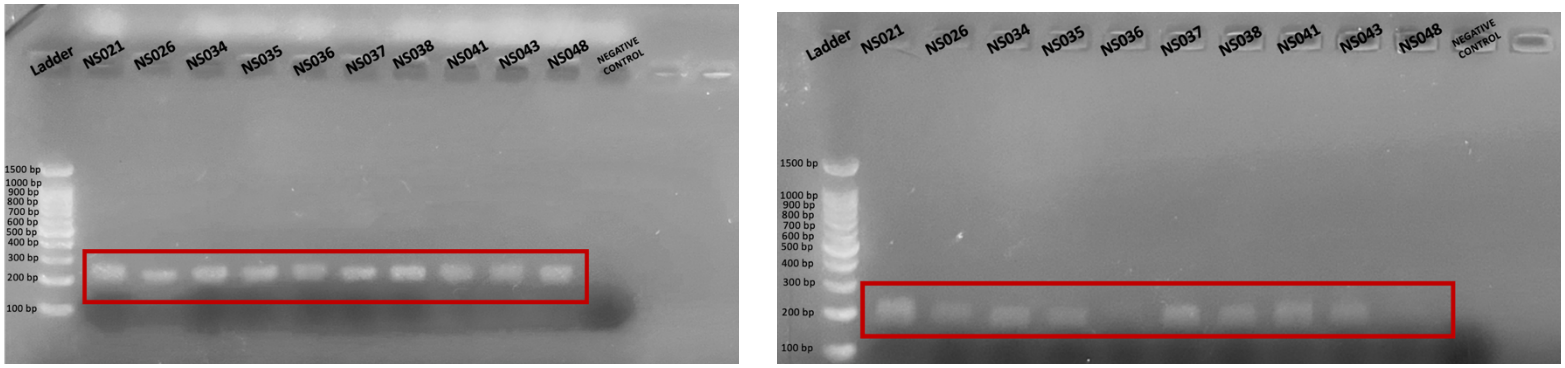


A B

Figure 6: Agarose gel electrophoresis of negative tick samples NS021, 026, 034, 035, 037, 038, 041, 043, 048. A) Synthesized cDNA from each sample underwent qPCR with *EHMT2* 6 primers, with an amplicon size of 207bp. B) Synthesized cDNA from each sample underwent qPCR with *EHMT2* 8 primers, with an amplicon size of 184bp
